# Meta-analysis shows both congruence and complementarity of DNA metabarcoding to traditional methods for biological community assessment

**DOI:** 10.1101/2021.06.29.450286

**Authors:** François Keck, Rosetta C. Blackman, Raphael Bossart, Jeanine Brantschen, Marjorie Couton, Samuel Hürlemann, Dominik Kirschner, Nadine Locher, Heng Zhang, Florian Altermatt

**Author notes:** François Keck and Rosetta Blackman should be considered joint first authors.

## Abstract

Assessment of the diversity and composition of biological communities is central to studies in ecology as well as for ecological monitoring. Historically, individual taxonomic groups have been assessed separately, while for an understanding of the state and change of biodiversity under ongoing global change an integrated assessment would be necessary. DNA metabarcoding has been proposed to be a highly promising approach especially for the assessment of aquatic communities, and numerous studies have investigated the consistency of this new technique with traditional morpho-taxonomic approaches. These individual studies have used DNA metabarcoding to assess diversity and community structure of aquatic organisms both in marine and freshwater systems globally over the last decade. However, a systematic analysis of the comparability and effectiveness of DNA-based community assessment across all of these studies has hitherto been lacking. Here we performed the first meta-analysis of all available studies comparing traditional methods and DNA metabarcoding to measure and assess biological diversity of key aquatic groups, including microorganisms, macroinvertebrates, and fish. Across 215 datasets, we found that DNA metabarcoding provides diversity estimates (richness) that are globally consistent to those obtained using traditional methods. DNA metabarcoding also generates species inventories that are highly congruent with traditional methods for fish. Contrastingly, however, species inventories of microorganisms and macroinvertebrates obtained by DNA metabarcoding showed pronounced differences to traditional methods, missing some taxa but at the same time detecting otherwise overseen diversity. Our results indicate that DNA metabarcoding is efficient to estimate local and regional richness. The method is generally sufficiently advanced to study the composition of fish communities and replace more invasive traditional methods. For smaller organisms, like macroinvertebrates and microorganisms, DNA metabarcoding may continue to give complementary rather than identical estimates compared to traditional approaches. Systematic and comparable data collection will increase the understanding of different aspects of this complementarity, and increase the effectiveness of the method and adequate interpretation of the results.

## Introduction

Assessment of biological assemblages is key to almost every study in ecology (Hampton et al., 2013), and managing and preserving ecosystems requires a global effort to regularly monitor the composition and diversity of their biological communities (Jørgensen, Xu, Salas, & Marques, 2010). In aquatic ecosystems, routine biological assessment has a long history and a wide range of groups of organisms (such as diatoms, aquatic plants, invertebrates, and fish) are used to monitor the state and change of aquatic environments over time, and to assess different types of human-induced pressures and impairments (Barbour, Gerritsen, Snyder, & Stribling, 1999; Borja, Franco, & Pérez, 2000; Hering et al., 2018). In order to use these organisms in large ecological monitoring programmes, a variety of methods have been developed and standardized based on the capture of individuals (such as biofilm collection, sampling by net, or electro-fishing), but ultimately the taxonomic identification is achieved using morphological criteria. This can be time-consuming, prone to errors, and lead to oversimplification, as taxa are often identified to a higher taxonomic level only (e.g., family or genus) (Haase et al., 2006; Mandelik, Roll, & Fleischer, 2010).

Technological advances in DNA sequencing (high throughput sequencing) and data analysis are currently revolutionizing biodiversity sciences, and are providing a novel approach to characterize biodiversity of whole communities by using DNA traces of organisms for their taxonomic identification (Hering et al., 2018; Leese et al., 2016). Thereby, metabarcoding can either be based on DNA extracted from bulk samples (i.e. sorted organisms) or from environmental samples (eDNA), with subsequent amplification of a specific gene region in target taxonomic groups using a dedicated primer pair (Pawlowski, Apothéloz-Perret-Gentil, & Altermatt, 2020; Taberlet, Coissac, Hajibabaei, & Rieseberg, 2012). For the purpose of this study we will refer to DNA metabarcoding of eDNA and bulk samples as DNA metabarcoding hereafter. DNA metabarcoding is widely seen as a promising solution to the constraints associated with traditional methods and is presented as being cheaper, faster, more sensitive, and easily scalable for routine monitoring programmes (Altermatt et al., 2020; Hering et al., 2018; Leese et al., 2016), and has been proposed to transform how animal and plant communities are surveyed (Deiner et al., 2017).

Along these lines, a growing number of articles have focused on testing the effectiveness of metabarcoding, and in particular its ability to quantify biodiversity and detect species present in the environment, over the last decade. Many studies compared the diversity of organisms assessed with DNA metabarcoding to assessments with traditional methods. Such comparisons are needed for two main reasons. First, to validate the concept and the protocols against methods which have been applied for decades and whose performance and limitations are well documented. Second, to ensure the continuity of long-term traditional monitoring time series. Accordingly, many case- or site-specific studies have attempted to estimate the congruence between the two approaches (e.g., Abad et al., 2016; Cahill et al., 2018; Fernández et al., 2018; Hänfling et al., 2016; Li et al., 2019; Mächler et al., 2019; Vasselon, Rimet, Tapolczai, & Bouchez, 2017). These studies have not only been conducted on a broad range of aquatic ecosystems, individual study sites and organismal groups, but are also reporting a wide range of comparability versus divergence to classically assessed community data. For example, individual studies often report that metabarcoding detects a substantial number of species identified on morphological criteria. However, the variability between studies is high, and often a significant fraction of diversity is still only detected by one or the other approach (e.g., Apothéloz-Perret-Gentil et al., 2017; Aylagas, Borja, Irigoien, & Rodríguez-Ezpeleta, 2016; Kelly et al., 2017; Polanco Fernández et al. 2021; Vasselon et al., 2017). Such knowledge on variability (or consistency) across studies is needed to informs about the specificities and limitations of each approach and to thus allows us to define the best strategies for current and future biomonitoring programs (Hering et al., 2018). To our knowledge, there has been no attempt to comprehensively review and analyse the available information in a systematic and quantitative way.

Here, we conducted the first systematic meta-analysis of the available studies that have compared the performances of DNA metabarcoding with traditional methods to estimate the diversity of a variety of organisms in aquatic ecosystems, both marine and freshwater. Specifically, we first studied which method detects the highest taxonomic diversity. Secondly, we studied if and to what extent these methods are congruent in the taxa detected. Finally, we examined the ecological scale (local versus regional) of this congruence and if the congruence between traditional methods and metabarcoding is dependent on the group of targeted organisms (microorganisms, macroinvertebrates, and fish). By aggregating the large body of literature available, we aim to end speculation as to the abilities of both methods for monitoring aquatic systems.

## Material and Methods

### Data collection

We conducted a systematic and comprehensive meta-analysis on all available, published studies following a set of formal criteria. Studies reporting comparisons between DNA metabarcoding and traditional methods to assess biological diversity in aquatic ecosystems (marine and freshwater) were searched using the online database Web of Science Core Collection (Clarivate Analytics) on the 25th February 2021. The combination of keywords used as search terms was chosen to be as specific as possible to include any studies comparing DNA metabarcoding to a traditional method (including both capture and identification) in aquatic ecosystems. Additionally, the search was limited to articles published between 2010 and 2021, since no study using metabarcoding on our targeted groups was expected prior to 2010 as the methods and apparatus needed were unavailable. The complete query used for the search is available as Supplementary Information (Supplementary Material S1). The initial search was complemented with a manual inspection of all articles published in *Metabarcoding and Metagenomics* and *Environmental DNA*, two new journals specialised in metabarcoding and environmental DNA (eDNA) studies, but whose publications were not yet indexed by Web of Science.

The initial search output was then carefully screened by manually checking the title, abstract and when necessary the complete content of the articles and their supplementary material, in order to retain only those articles that met our inclusion criteria. To be included in our analysis, studies had to report a comparison between DNA metabarcoding (on bulk or eDNA samples) and one or several traditional methods to assess biological diversity in aquatic ecosystems (marine or freshwater). The diversity estimated by the two approaches had to be reported using quantitative values expressed as taxonomic richness (i.e. the number of detected taxa). When this information was not directly available, but authors provided taxonomic lists obtained with metabarcoding and traditional methods, we were able to estimate the values of diversity needed for the analyses. We included only studies using DNA metabarcoding to assess taxonomic diversity at the community level and therefore excluded studies using conventional PCR, qPCR, ddPCR, or barcoding for single species detection. Studies using artificially assembled communities (e.g., mock communities, aquarium) were also excluded. Finally, we excluded studies focusing on bacteria and fungi for which communities are rarely assessed using traditional methods.

### Data extraction

For each comparison between a traditional method and DNA metabarcoding, the measures of diversity (richness) were extracted from the articles and the supplementary materials published by the authors at two different levels: local alpha diversity (i.e. the average diversity per site) and regional gamma diversity (i.e. the total diversity across sites).

When taxonomic diversity (here richness) is assessed with two different approaches (here traditional and DNA metabarcoding), the total number of detected taxa can be decomposed into three subsets: the subset of taxa detected only by the traditional method, the subset of taxa detected only by DNA metabarcoding and the subset of taxa detected by both methods, i.e. the intersection subset (Figure 1).

**Figure 1.**
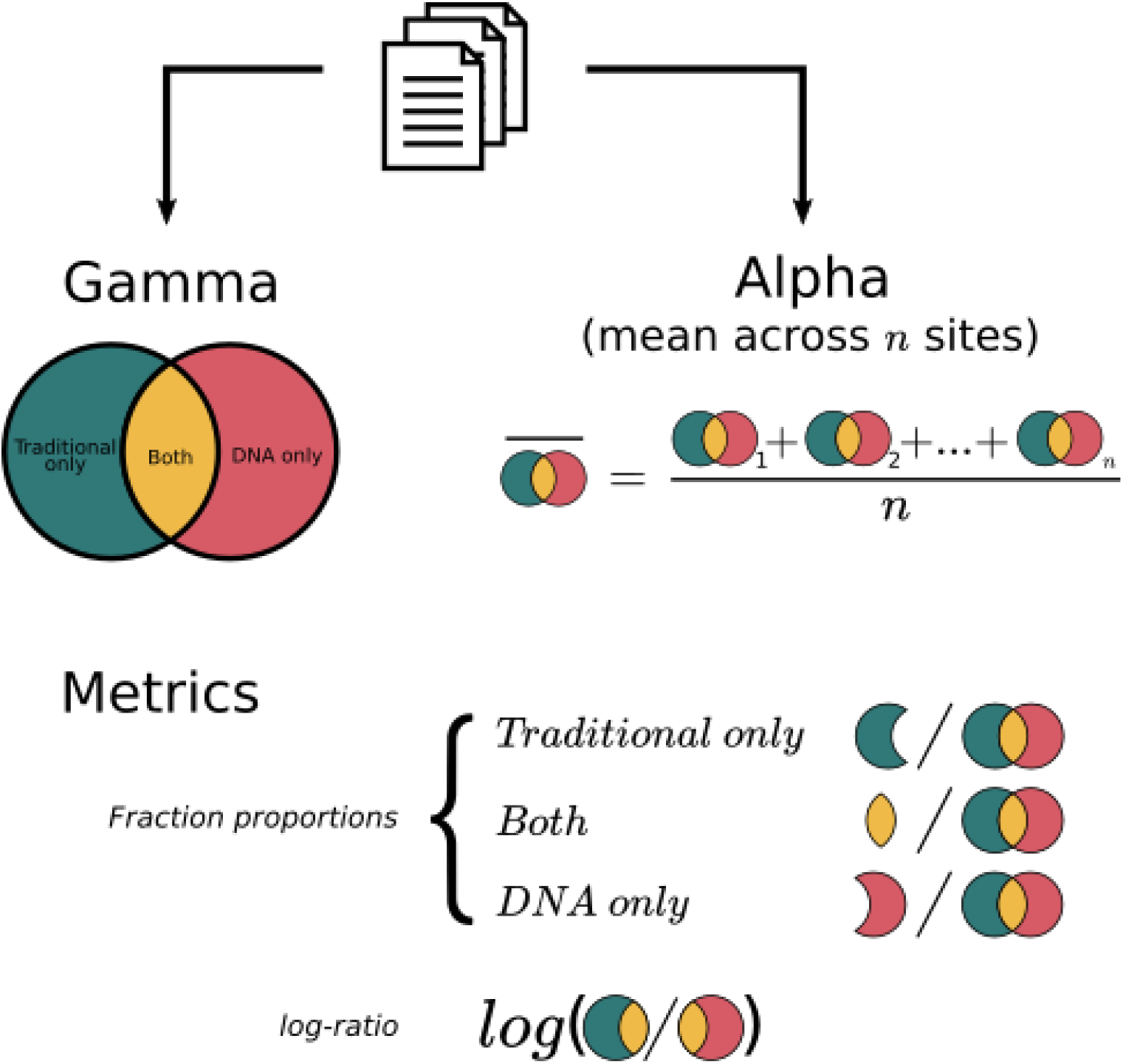
Study workflow. Gamma diversity (i.e., regional richness) and alpha diversity (i.e., local richness) values were extracted for different taxonomic groups from 99 studies. For each type of diversity, the relative fraction of taxa detected by the traditional method only (green), by DNA metabarcoding only (red) and by both methods (yellow) were compared. The log-ratio between the total diversity detected by the traditional method and the total diversity detected by DNA metabarcoding was also assessed.

For gamma diversity, measures of these three subsets are often reported in publications in the main text or in figures (e.g., Venn diagrams or stacked bar plots) and were directly extracted for each comparison. In contrast, the subsets of taxa detected only by one or the other method are rarely reported at the level/resolution of each site, but are often only available in an integrated manner only. Therefore, for each method, we computed an average of alpha diversity (local richness) across all sites. We did the same for the intersection fraction (i.e. the subset of taxa detected by both methods). The intersection fraction for alpha diversity is rarely reported, but could be estimated when the authors provided detailed lists of taxa detected by the two methods. Finally, we estimated the traditional only fraction and the DNA only fraction by subtracting the intersection fraction from the traditional and DNA total alpha diversity.

Along with the measures of diversity, we also extracted key variables of interest including article metadata, information on the study designs (sampled habitat, targeted taxonomic group, taxonomic resolution) and methodological details about the metabarcoding approach (markers, primers, technologies). The complete list of extracted data is provided as Supplementary Table S1. To simplify the analyses and ease the reading of the results, the organisms targeted in the included studies were grouped into four categories: microorganisms (including diatoms, zooplankton, phytoplankton, cyanobacteria, protists), macroinvertebrates, fish, and “others” (e.g., corals, macrophytes, amphibians); the latter only containing a small number of studies.

### Statistical analyses

To assess whether one approach detected more diversity (richness) than the other, we used the log-ratio *ln(A/B)* where, for each method comparison, *A* is the total diversity detected by the traditional method and *B* is the total diversity detected by DNA metabarcoding (Figure 1). The log-ratio is a widely used effect size measure in ecological meta-analyses to summarize the magnitude and direction of multiple research outcomes (Hedges, Gurevitch, & Curtis, 1999). When *A* and *B* are equal (i.e. the two methods estimate the same diversity) the log-ratio value is zero. To test which method detected the highest diversity we used linear mixed models with the study block as random effect (intercept). Two models were fitted: an intercept-only model to address the mean log-ratio and one including the group of organisms as an independent variable to test possible effects of the type of taxa on the log-ratio.

To analyse the congruence between traditional methods and DNA metabarcoding, we compared the three fractions (traditional only, DNA metabarcoding only and the intersection fraction, see above and Figure 1). Fractions were standardized by turning them into proportions with values ranging from zero to one to allow cross-study comparisons. To test differences between fractions within and across groups of taxa, we used a beta-regression mixed model (Ferrari & Cribari-Neto, 2004) including the taxonomic groups (microorganisms, macroinvertebrates, and fish) and the types of fraction (traditional only, DNA only, and both) as independent variables. Two models were fitted separately for gamma and alpha diversity and two levels of nested random effects (intercepts) were specified (studies and comparisons) for these models. Proportion values were compressed using the method of Smithson and Verkuilen’s (2006) to avoid true zeros and ones in the beta-regression. Due to a strong lack of data in the other taxonomic levels and large class imbalance, the model was fitted only for the comparisons made at species level (i.e. the level for which we had the most data available). Beta-regression models were completed with post hoc pairwise comparisons among groups based on estimated marginal means with a Tukey procedure to control for family-wise error rate.

We performed all the statistical analyses with R 4.0.3 software (R Core Team, 2020). Generalized linear mixed models were fitted using the glmmTMB package (Brooks et al., 2017) and the emmeans package was used to perform post hoc pairwise comparisons (Lenth, 2021).

## Results

From the 1,217 papers initially identified by the literature search, many focused on single species assays or bacteria. In total, 99 studies met our inclusion criteria and were used in the analyses (Table 1, Supplementary Material S2). From these, we extracted a total number of 215 comparisons (some articles presented several comparisons using different primers or different taxonomic groups assessed separately) of diversity measurements between a traditional method and DNA metabarcoding. Most of these comparisons were presented at regional level, that is, gamma diversity (188 comparisons). A relatively large number of studies reported data for each approach at site level (120 comparisons), while the intersection fraction between the traditional and DNA methods could be extracted for 88 comparisons.

**Table 1.** List of articles included in the meta-analysis sorted by group of organisms.

| Group of organisms | Articles included |
| --- | --- |
| Microorganisms | Abad et al. (2016), Apothéloz-Perret-Gentil et al. (2017), Apothéloz-Perret-Gentil et al. (2020), Bachy et al. (2013), Bailet et al. (2020), Clarke et al. (2017), Djurhuus et al. (2018), Dzhembekova et al. (2020), Eiler et al. (2013), Harvey et al. (2017), Hirai et al. (2015), Huang et al. (2020), Huo et al. (2020), Kang et al. (2021), Kermarrec et al. (2014), Kim et al. (2019), Kim et al. (2020), Li et al. (2019), Liu et al. (2017), Minerovic et al. (2020), Mora et al. (2019), Nunes et al. (2019), Pérez-Burillo et al. (2020), Pujari et al. (2019), Rivera et al. (2018), Rivera et al. (2018), Schroeder et al. (2020), Semmouri et al. (2021), Vasselon et al. (2017), Visco et al. (2015), Xiao et al. (2014), Yang et al. (2017), Zimmermann et al. (2015) |
| Macroinvertebrates | Aylagas et al. (2016), Aylagas et al. (2018), Azevedo et al. (2020), Borrell et al. (2017), Cahill et al. (2018), Cowart et al. (2015), Elbrecht et al. (2017), Emilson et al. (2017), Erdozain et al. (2019), Fernández et al. (2018), Fernández et al. (2019), Haenel et al. (2017), Harper et al. (2020), Kelly et al. (2017), Krol et al. (2019), Kuntke et al. (2020), Laini et al. (2020), Leese et al. (2021), Lejzerowicz et al. (2015), Lobo et al. (2017), Mächler et al. (2019), Marshall et al. (2020), Martins et al. (2019), Obst et al. (2020), Rivera et al. (2021), Serrana et al. (2018), Serrana et al. (2019), Steyaert et al. (2020), Sun et al. (2019), Uchida et al. (2020), Vivien et al. (2019) |
| Fish | Afzali et al. (2021), Aglieri et al. (2020), Berger et al. (2020), Bleijswijk et al. (2020), Boivin- Delisle et al. (2021), Bylemans et al. (2018), Cilleros et al. (2019), Closek et al. (2019), Collins et al. (2019), Doble et al. (2020), Evans et al. (2017), Polanco Fernández et al. (2021), Fujii et al. (2019), Goutte et al. (2020), Hänfling et al. (2016), Hayami et al. (2020), McClenaghan et al. (2020), McDevitt et al. (2019), Nguyen et al. (2020), Oka et al. (2021), Olds et al. (2016), Pont et al. (2018), Port et al. (2016), Sakata et al. (2021), Sard et al. (2019), Shaw et al. (2016), Snyder et al. (2020), Thomsen et al. (2012), Valentini et al. (2016) |
| Others<br>(including multiple<br>groups) | Alsos et al. (2018), Deagle et al. (2018), Gran-Stadniczenko et al. (2017), Leduc et al. (2019), Nichols et al. (2019), Shackleton et al. (2019), Valentini et al. (2016) |

The included several continents and climatic regions. However, data are strongly spatially aggregated and most studies are focused on Europe and North America (Figure 2 A). The dataset covers a large variety of functional and taxonomic groups that range from microbial species to fish (Figure 2 B), representing an important variation in body size or trophic position. Both marine (44% of comparisons) and freshwater ecosystems (56%) are represented (Figure 2 C), with data available for numerous types of aquatic environments, ranging from small streams and ponds to oceans (Supplementary Figure S1). Most comparisons were made and reported by the authors at species level (49.8% of comparisons), although a significant number of studies investigated diversity at higher taxonomic levels (Figure 2 D). The variety of studied organisms and habitats is reflected by the diversity of genetic markers (9 markers, 64 different primer pairs) and source of DNA used by the authors (Figure 2 E-F). Finally, several sequencing technologies are represented in the dataset, with Illumina MiSeq being the most commonly used technology over the studied period (69.3% of comparisons; Figure 2 G). Detailed distributions of the recorded observations are shown in Supplementary Figure S1 and S2.

**Figure 2.**
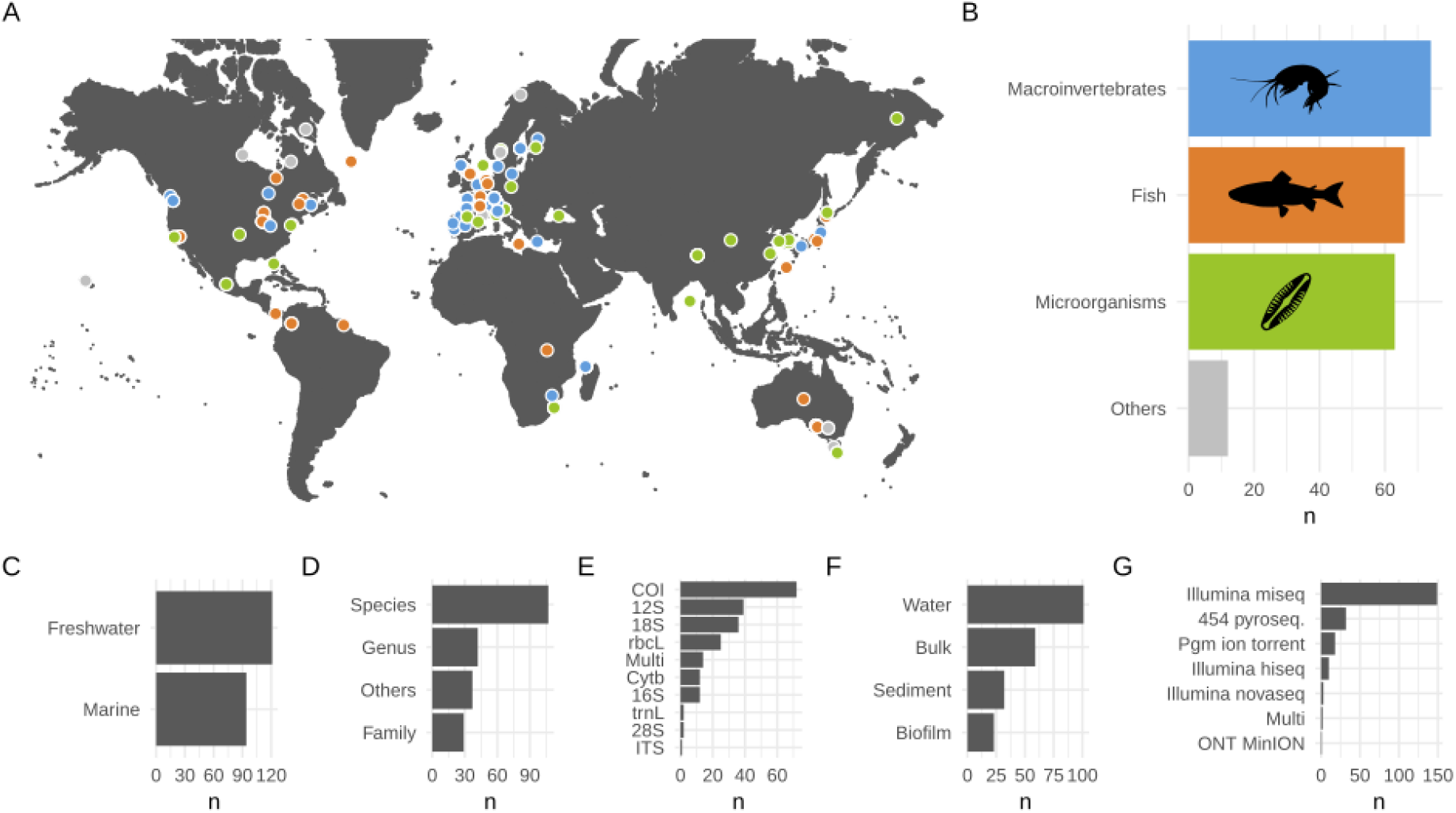
Overview of the different comparisons extracted from the studies included in the meta-analysis. **A.** Geographic location of the comparisons. Colors indicate the group of organisms used (microorganisms in green, macroinvertebrates in blue, fish in orange and other types in light grey). **B.** Number of comparisons (n) across the different groups of organisms. **C-G.** Number of comparisons (n) across biomes (marine includes brackish waters), taxonomic levels of identification, genetic markers, origins of DNA and sequencing technologies. The “multi” category includes comparisons combining several other categories.

We measured an average log-ratio between the traditional and DNA metabarcoding of - 0.010 (sd = 0.764) for gamma diversity (Figure 3). This average log-ratio was not found to be significantly different from zero by the statistical model (Z-value = −0.187, p-value = 0.851) and the effect of the group of organisms was not found significant (Wald Chi2 = 2.078, df = 3, p-value = 0.556). For alpha diversity, the average log-ratio was 0.093 (sd = 0.73) and was found marginally-significant in the model (Z-value = 1.979, p-value = 0.048). Similarly to the gamma diversity, the effect of the group of organisms was not found significant for alpha diversity (Wald Chi2 = 0.486, df = 3, p-value = 0.922). Details on the estimated terms of the models are provided as supplementary information (Supplementary Table S2).

**Figure 3.**
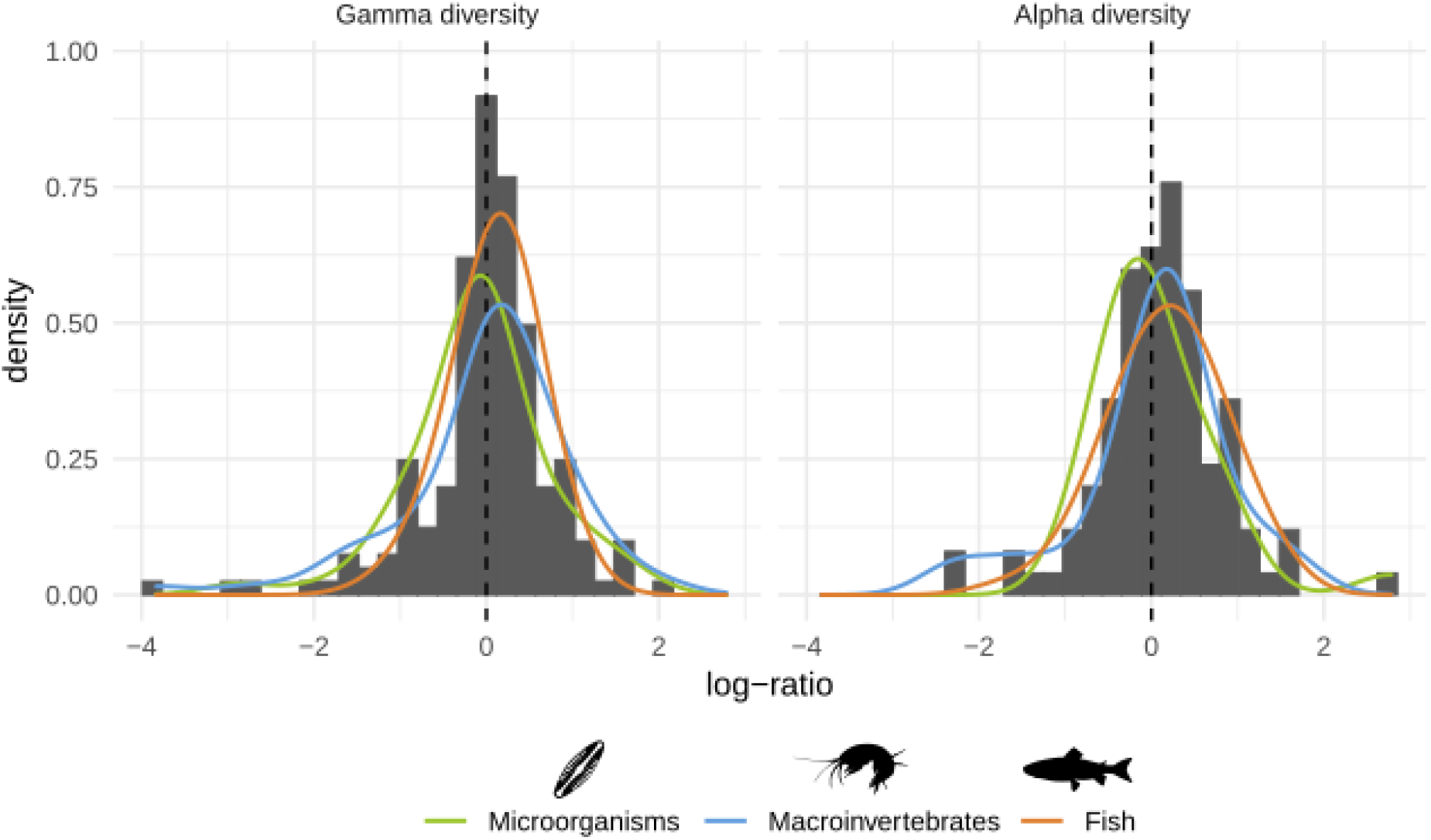
Histograms of the log-ratio between the total diversity detected by the traditional method and the total diversity detected by DNA metabarcoding. The left panel shows gamma diversity (i.e. regional richness) and the right panel alpha diversity (i.e. mean local richness). Density estimates (kernel bandwidth = 0.25) for each group of organisms are represented as colored overlays.

Overall, we found that the proportion of diversity detected was strongly dependent on the type of fraction considered (traditional only, DNA only, or both methods) and the group of organisms (Figure 4 for species, see Supplementary Figures S3 and S4 for the other taxonomic levels). This was confirmed both for alpha and gamma diversity by the analyses of deviance of the beta-regression models (interaction term p-values = 0.007 and <0.001 respectively, Supplementary Table S3). Further, we found that the fractions of gamma diversity detected only by the traditional method and only by DNA metabarcoding were significantly lower than the fraction detected by both methods for fish (Supplementary Table S4). This trend is inverted for the microorganisms (Figure 4 A), although only the “traditional-only” fraction was significantly higher than the “both” fraction (Supplementary Table S4). Macroinvertebrates are intermediate, with none of the fraction being significantly higher than the others (Figure 4 B and Supplementary Table S4). Very similar trends were observed for alpha diversity (Figure 4 D-F), but none of the pairwise tests showed significant differences (Supplementary Table S5), likely because of a lack of statistical power.

**Figure 4.**
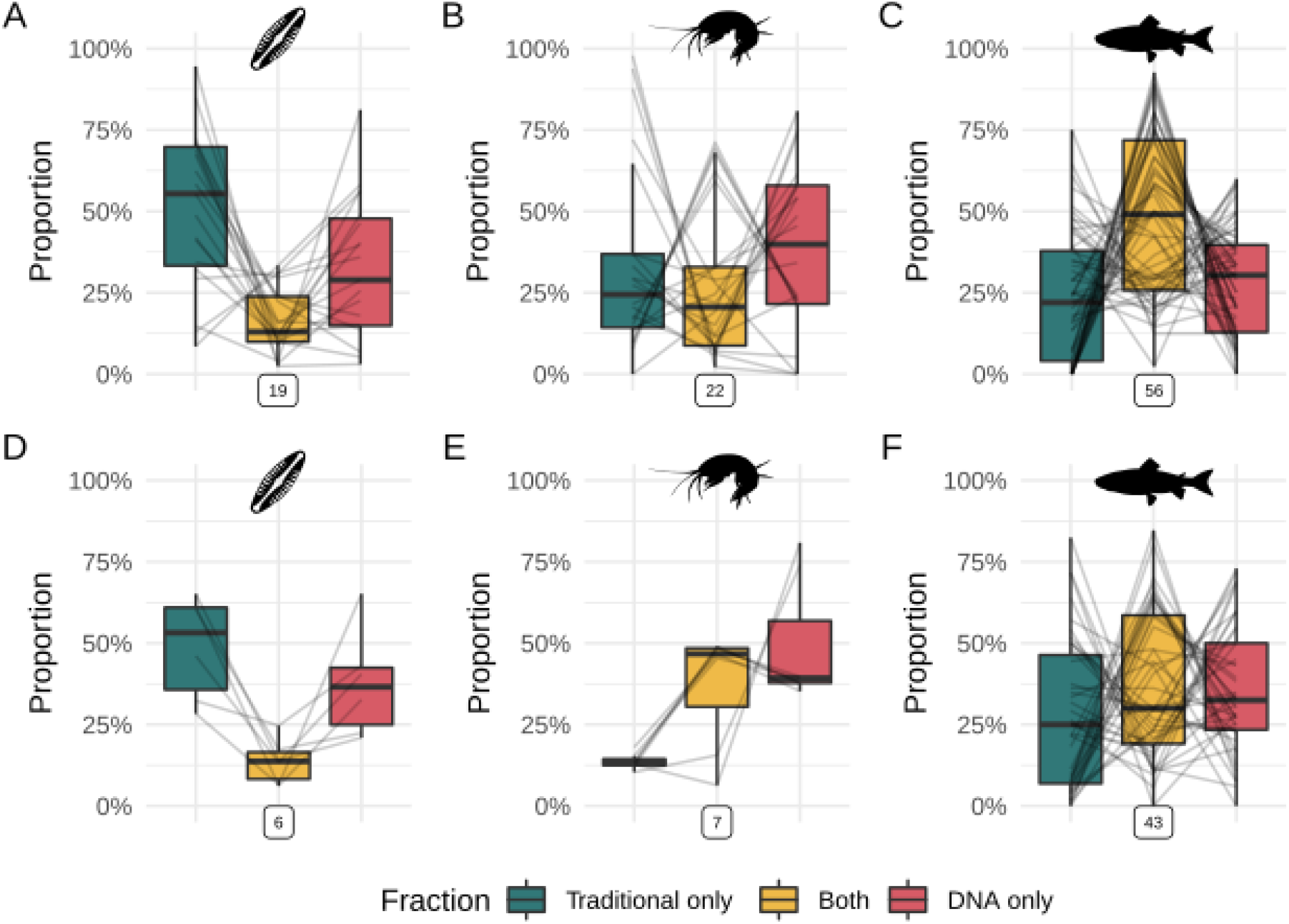
Relative fraction of diversity detected by the traditional method only, by DNA metabarcoding only and by both methods. Data are presented for different groups of organisms identified at species level only. Boxplots show medians, first and third quartiles, and full ranges (limited to 1.5 × interquartile range). Grey lines connect values from the same comparison. Framed numbers below each panel indicate the number of comparisons represented. **A-C.** Gamma diversity for microorganisms, macroinvertebrates and fish. **D-F.** Alpha diversity for microorganisms, macroinvertebrates and fish.

We also observed a high variability in the proportion of species found with both methods, ranging from 2% (macroinvertebrates) to 93% (fish). This variability was also important within taxonomic groups (e.g., ranging from 2% to 71% in macroinvertebrates, Figure 4).

Besides the group of organisms, we also recorded and explored the effects of other factors like the origin of DNA or the difference between marine and freshwater systems. The effects of these factors and especially the combined effects of these factors with the effect of the group of organisms are only visually presented (Supplementary Figures S5–S8) but were not formally tested because of a lack of data, class imbalance, and the high multicollinearity among variables (e.g., between the groups of organisms and the origin of DNA).

## Discussion

Since the initial proposal to use metabarcoding to simultaneously identify all organisms in the environment (Taberlet et al., 2012), many scientists are intrigued by or have questioned the comparability of this new approach with the traditional methods used for biodiversity inventories. In order to synthesise and provide a quantitative basis of the current state of knowledge on this issue, we collected and analysed all available data from a large corpus of publications comparing traditional methods with DNA metabarcoding.

At regional level (gamma diversity), we found that there was no one approach that detected more diversity (richness) than the other. This result indicates that the regional richness estimated by metabarcoding matches well with the traditional methods, hence confirming the potential of this approach for biodiversity assessment in aquatic ecosystems. This is, however, an overall trend across all studies, while the distribution of log-ratio (Figure 3 A) shows a wide range of situations, with some rare cases where the absolute log-ratio reaches 2 (i.e. one method detected 7 times more taxa at the regional scale than the other). Unlike gamma diversity, the log-ratio was found significantly higher than zero for alpha diversity suggesting that at site level, DNA metabarcoding detects more diversity than its traditional counterparts. This result should be interpreted with caution, especially in view of the small effect size reported (on average metabarcoding detected 1.1 times more taxa than the traditional methods), but is in line with a recent meta-analysis showing that the probability of species detection is higher with eDNA than with traditional methods (Fediajevaite, Priestley, Arnold, & Savolainen, 2021).

Traditional methods and their respective efficacy will likely remain stable in the future as those methods have been established, optimised and standardized over decades. Contrastingly, the techniques behind DNA metabarcoding are continuously refined and the approach has likely not yet reached its full potential in its ability to detect taxonomic diversity (Keck, Vasselon, Tapolczai, Rimet, & Bouchez, 2017). In particular, progress in sequencing technologies, bioinformatics processing, and reference database coverage will improve the capacity of DNA metabarcoding to estimate diversity. This margin of progress may suggest that metabarcoding could widen the gap with traditional approaches in terms of its ability to measure taxonomic richness in the future.

Our analyses did not demonstrate a significant effect of the type of organism on the log-ratio between the number of taxa detected by DNA and traditional methods. This is surprising and a bit unexpected, as especially for macroinvertebrates there have been extensive debates in the literature about the suitability of (e)DNA metabarcoding (Blackman et al., 2019; Elbrecht et al. 2017). The large variation observed in the log-ratio across studies, however, suggests that the accuracy and overlap depends also on the combination of many other factors, which can not be modelled here due to a lack of data. Besides the group of organisms, the context of the study and the methods used can also have drastic effects on the relative estimates of biodiversity provided by DNA metabarcoding and traditional methods. For instance, eDNA in lentic environments may persist longer than in lotic or marine systems (Collins et al., 2018; Dejean et al., 2011), which implies that metabarcoding data might have a better congruence with traditional methods. Whereas eDNA from lotic and marine systems is influenced by transportation from flow (Deiner & Altermatt, 2014) and ocean currents (Harrison, Sunday, & Rogers, 2019), meaning that the DNA does not represent the same scale as a traditional sample from the same environment. For example, a biofilm DNA sample from a small pond and a water DNA sample from the deep ocean are not representing the same scales. Therefore, the eDNA samples will reflect a very different scale both in time and space (Civade et al., 2016; Deiner, Fronhofer, Mächler, Walser, & Altermatt, 2016) compared to traditional point sampling, such as kick-nets (Mächler et al., 2019). Thus, the large diversity of habitats and sampling strategies in our dataset can possibly explain the large variation in log-ratio values.

Although DNA and traditional methods estimated the same number of taxa (diversity) on average, our results suggest that they often do not count the same taxa. This is shown by the fraction of species detected by both methods which is particularly low for microorganisms and macroinvertebrates. This result has important implications, because in addition to taxonomic richness, taxonomic composition is an essential element of biodiversity. Ecologists are interested in species identities in biological assemblages because they are rich in information about environmental quality and ecosystem functioning. Additionally, being able to identify taxa is important to monitor rare and endangered species or to detect invasive species. It is worth noting that, in this study, the correspondence between metabarcoding and traditional taxonomic lists is only based only on the exact match between taxa names. The taxonomic and phylogenetic distances between the methods are likely to be less significant, which can be used to optimize the calculation of ecological quality indices (Keck, Vasselon, Rimet, Bouchez, & Kahlert, 2018). Such metrics, however, are rarely reported and could not be evaluated.

For microorganisms, which are mainly represented by benthic diatoms and planktonic protists, the discrepancies between traditional methods and DNA metabarcoding can be attributed to the respective flaws of these two approaches, which are well documented. Traditional approaches that rely on microscopic morphological characters for species identification are known to be a source of errors. In particular, several species and species complexes which have been initially separated using molecular methods are difficult to identify on the sole basis of light microscopy (Jahn et al., 2017; Pinseel et al., 2017). Contrastingly, the DNA metabarcoding approach is also limited in recovering diversity unveiled by traditional methods. Firstly, the short DNA fragment used as a genetic barcode can be insufficient (in size and/or variability) to separate morpho-taxonomic species (Apothéloz-Perret-Gentil et al., 2017). Secondly, reference databases are still incomplete in terms of the diversity found in microbial communities, and many environmental sequences cannot be accurately classified at species level (Lindeque, Parry, Harmer, Somerfield, & Atkinson, 2013; Weigand et al., 2019). In this respect, coordinated efforts to build extensive reference databases for protists and other microbial species (e.g., Guillou et al., 2013; Rimet et al., 2019) are particularly important and are expected to improve metabarcoding performances in the future.

Like for microorganisms, the fraction of macroinvertebrate species detected by both methods (i.e., overlap in identity of organisms detected) is remarkably low. Again, the discrepancies between the lists of taxa produced using DNA metabarcoding and traditional morphological analysis can be explained by the inherent biases of both methods, as described above (Lobo, Shokralla, Costa, Hajibabaei, & Costa, 2017). Furthermore, it is important to mention that the taxonomic extent targeted by the two methods can be very different. The generic primers used for macroinvertebrate metabarcoding are often rather unspecific and target a diversity of organisms (based on their phylogenetic and thus primer-binding site similarity), which is often overlooked by the operators performing traditional identification (Elbrecht, Vamos, Meissner, Aroviita, & Leese, 2017). The recent development of specific primers for macroinvertebrate communities could help address this issue (Leese et al., 2021), but likely will not completely resolve it, as a polyphyletic group such as “macroinvertebrates”, solely defined by their size, may never be adequately captured by genetic methods without also including closely related but smaller organisms. Additionally, the large variance observed in the fraction detected by both methods can be explained by the fact that our dataset includes both environmental DNA and bulk samples for macroinvertebrates. Recent studies have shown that environmental DNA includes a larger diversity of taxa than bulk samples, but many are non-targeted and do not match the taxa identified by traditional methods (Gleason, Elbrecht, Braukmann, Hanner, & Cottenie, 2020; Macher et al., 2018).

For fish, the observed trend is clearly reversed to what we see with microorganisms and macroinvertebrates. The fraction of fish species detected by both methods indicates that the concordance is good between DNA metabarcoding and the traditional approaches. This result is in line with conclusions from individual studies (e.g., Valentini et al., 2016; Hänfling et al., 2016; Pont et al., 2018; Li et al., 2019). The good match of taxonomic lists generated by the two approaches can be explained by the limited regional diversity of fish communities investigated in the available studies. Compared to microorganisms and macroinvertebrates, regional fish faunas are often well documented, and reference databases are extensive and can be easily completed by sequencing new individuals (Valentini et al., 2016), Moreover, fish being not as genetically divergent as microorganisms and macroinvertebrates, scientists have been able to develop highly specific primers for this group (Kelly, Port, Yamahara, & Crowder, 2014; Miya et al., 2015; Valentini et al., 2016) which match well with the diversity of organisms captured and identified using traditional methods.

It is important to note that our conclusions are drawn using one standard measure of diversity, namely richness. Other metrics, like Shannon or Simpson indices that take into account of the relative abundance of taxa, could lead to different conclusions, in particular because the issue of abundance with metabarcoding is yet to be solved (Piñol, Mir, Gomez-Polo, & Agustí, 2015; Visco et al., 2015). Moreover, we did not investigate how comparable beta diversity (i.e. the species turnover among sites) was between the two methods. This is because beta diversity is more complicated to record and synthesize from multiple sources, as it is often reported using different metrics, also calling for more standardized reporting of data enabling future assessments and meta-analyses. Nonetheless, beta diversity remains an important component of diversity and can give different results if analysed through DNA or traditional methods (Bleijswijk et al., 2020).

Recently, several meta-analyses have investigated a variety of questions regarding eDNA and DNA metabarcoding (Duarte, Leite, Feio, Costa, & Filipe, 2021; Fediajevaite et al., 2021; Yates, Fraser, & Derry, 2019). This indicates that the field is gaining maturity and there is now sufficient data available to synthesize and draw general conclusions. It must be noted that this effort is possible only if studies report relevant and extractable data in comparable, complete, and standardized ways. Thus, we stress the importance that future studies comparing traditional methods and DNA metabarcoding provide consistent information about methods and results. As a starting point, we suggest that all variables listed in Supplementary Table S1 should be systematically reported in an accessible way. Adhering to simple standards will help to improve reproducibility and comparability among studies and facilitate future syntheses (Gerstner et al., 2017).

In conclusion, while DNA metabarcoding has great potential for biodiversity assessment in aquatic ecosystems, we need to consider the implications of significant discrepancies between traditional methods and DNA metabarcoding-based methods for particular organismal groups. Here, we showed that DNA metabarcoding and traditional methods give similar estimates of taxonomic richness across major organismal groups, which can make these tools interoperable for example, to study patterns and trajectory of biodiversity at large spatial and temporal scales. Importantly, however, the two approaches often still differ on the identity of the species detected, especially in macroinvertebrate and microorganism communities, yet give similar numbers of total taxa recorded. This may be a problem if the objective is to replace one method with another in long-term monitoring programs where taxon identity is important. Our results suggest that DNA metabarcoding is ready to replace invasive methods traditionally used to study fish communities, while for smaller organisms like macroinvertebrates and microorganisms, it will be a complementary approach, capable of revealing aspects of biodiversity that were previously ignored or underestimated by traditional methods.

## Acknowledgements

Funding is from the Swiss National Science Foundation Grants No PP00P3_179089 and 31003A_173074, the Swiss Federal Office for the Environment (FOEN/BAFU), and the University of Zurich Research Priority Program “URPP Global Change and Biodiversity” (all to F.A.). The amphipod silhouette picture was kindly provided by Roman Alter.

## Data and Code Accessibility

The data and R scripts to reproduce the analyses and results are available at: [availability upon publication].

## Author Contributions

FA, FK, and RCB conceived the study. All authors collected the data. FK analysed the data, FK, RCB, and FA wrote the paper with contributions from all authors.

