## Supplementary Information for "Meta-analysis shows both congruence and complementarity of DNA metabarcoding to traditional methods for biological community assessment"

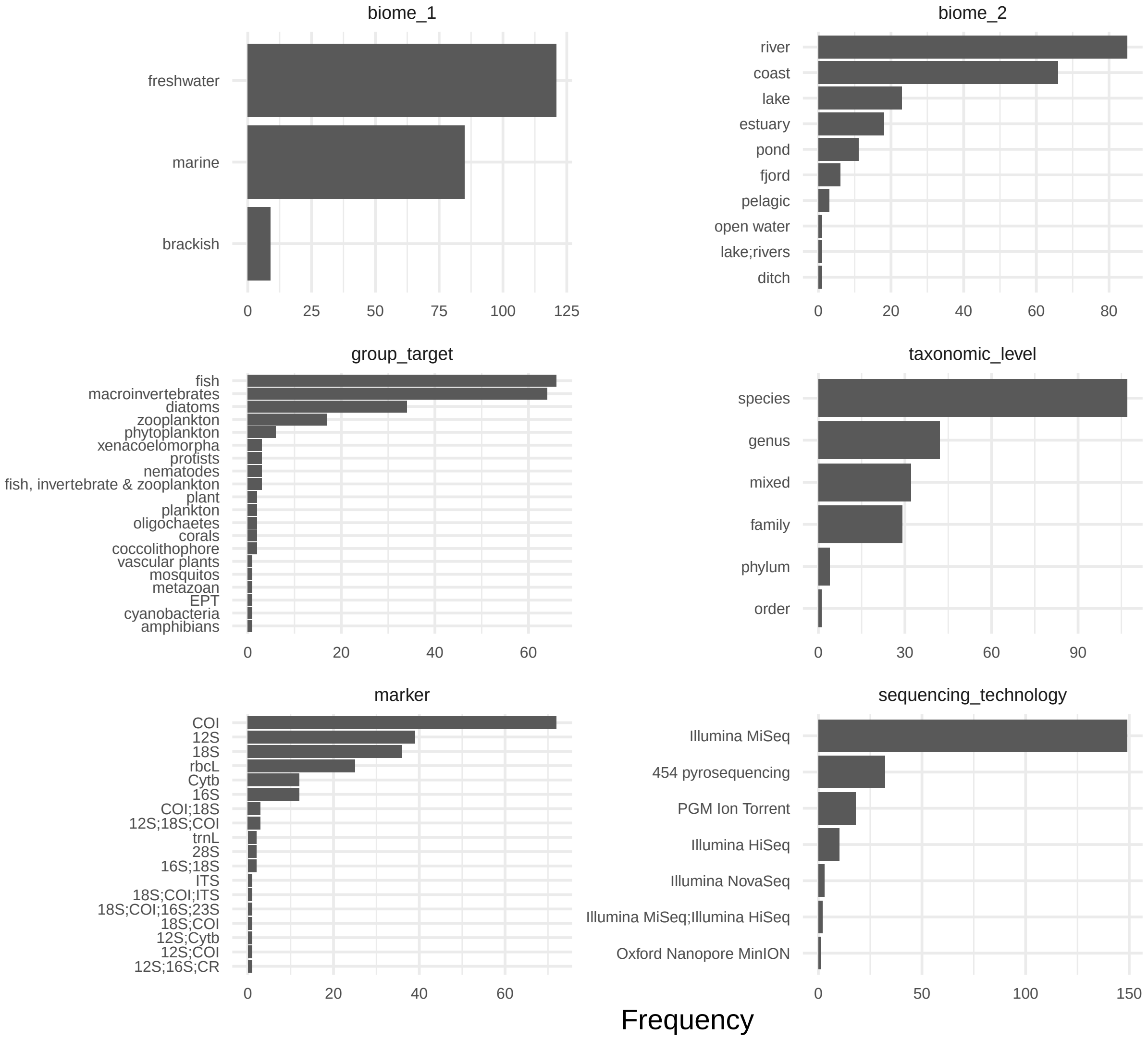

**Supplementary Figure 1.** Number of observations across the different categories for each categorical variable recorded. Categories are represented as they were recorded and before the simplification used in Figure 2.

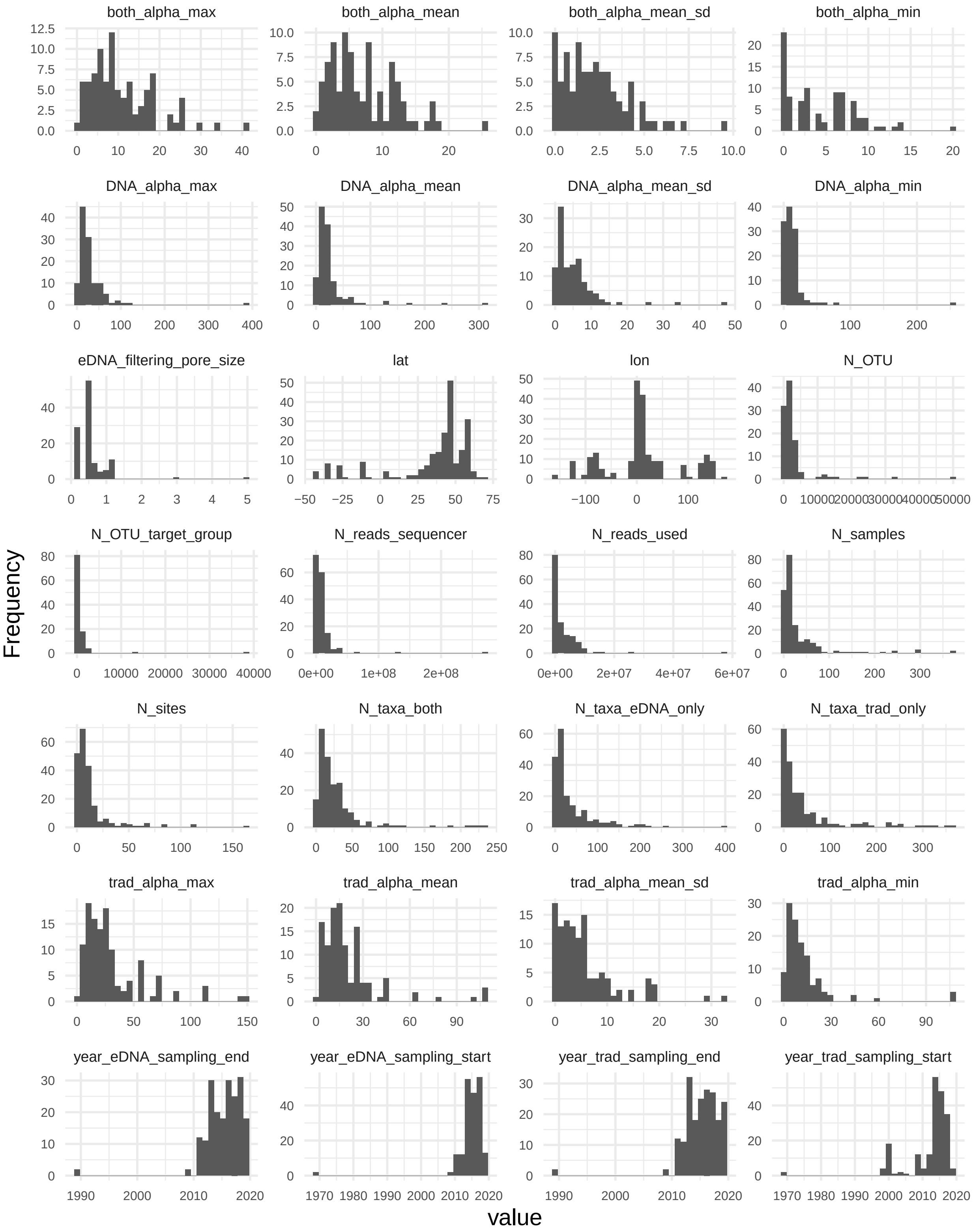

**Supplementary Figure 2.** Histograms of observations for each continuous variable recorded.

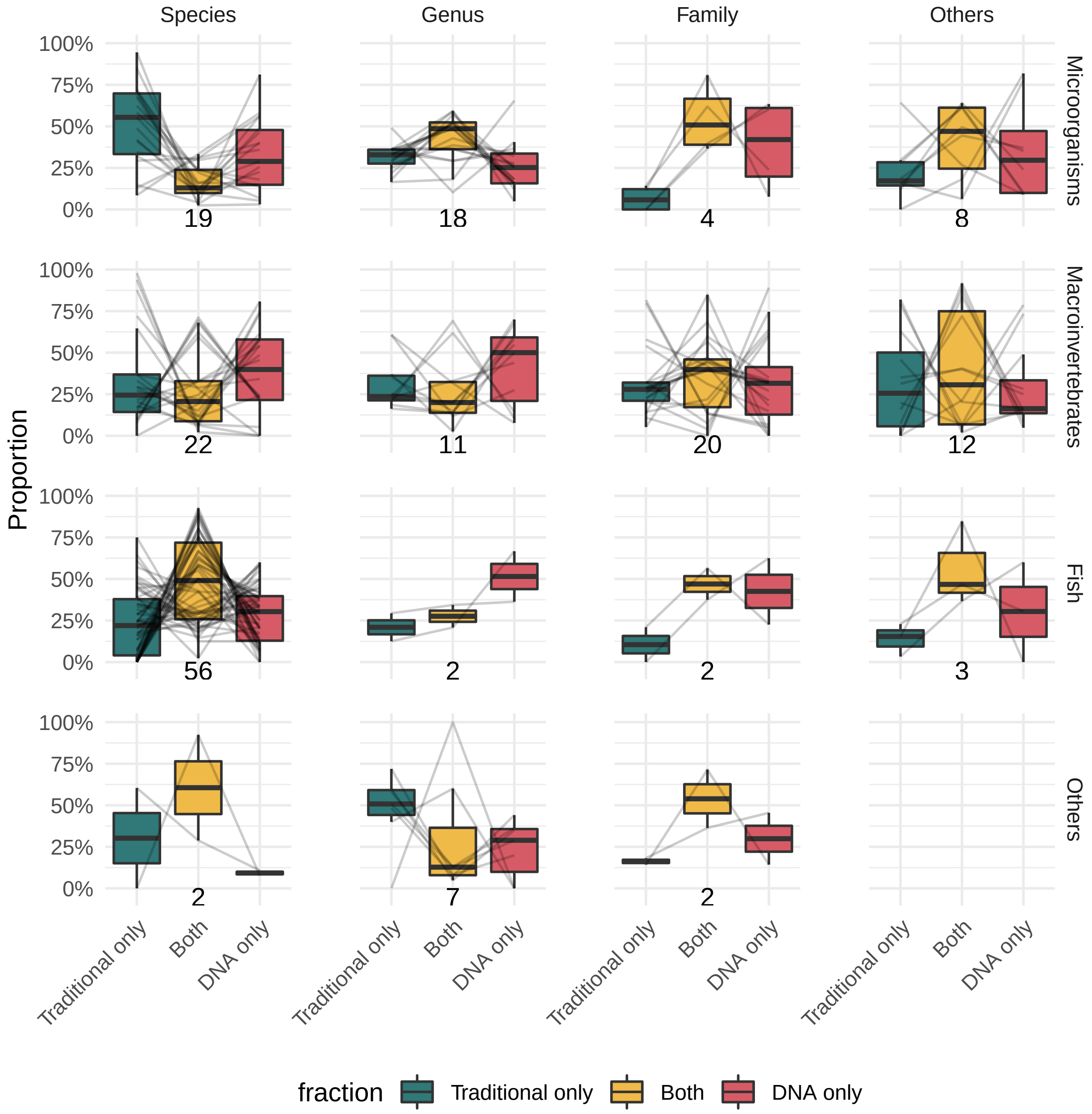

**Supplementary Figure 3.** Relative fractions of gamma diversity detected by the traditional method only, by DNA metabarcoding only and by both methods. Data are presented for different groups of organisms identified at different taxonomic levels. Boxplots show medians, first and third quartiles, and full ranges (limited to 1.5 × interquartile range). Grey lines connect values from the same comparison. Numbers below each panel indicate the number of comparisons represented.

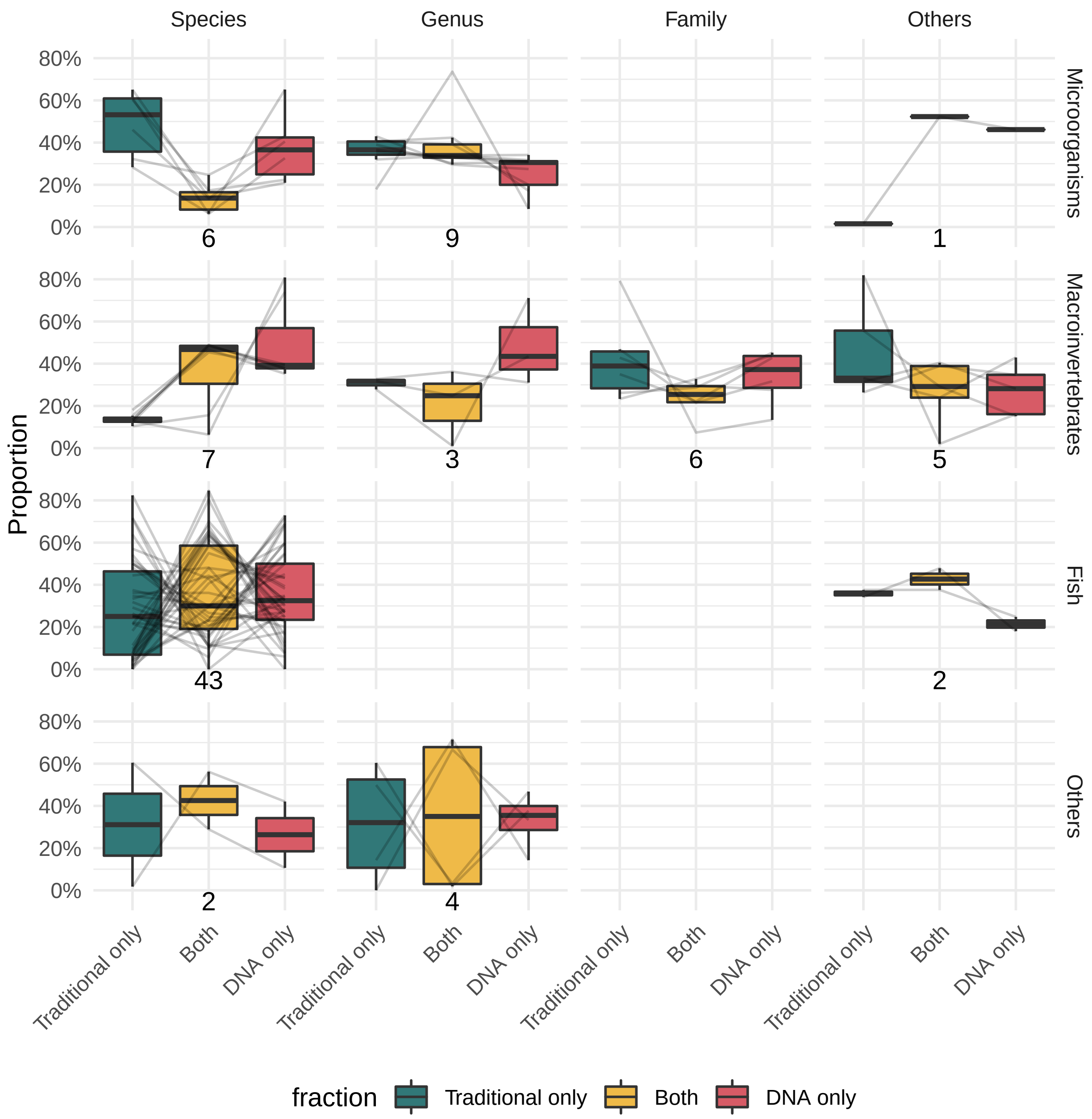

**Supplementary Figure 4.** Relative fractions of alpha diversity detected by the traditional method only, by DNA metabarcoding only and by both methods. Data are presented for different groups of organisms identified at different taxonomic levels. Boxplots show medians, first and third quartiles, and full ranges (limited to 1.5 × interquartile range). Grey lines connect values from the same comparison. Numbers below each panel indicate the number of comparisons represented.

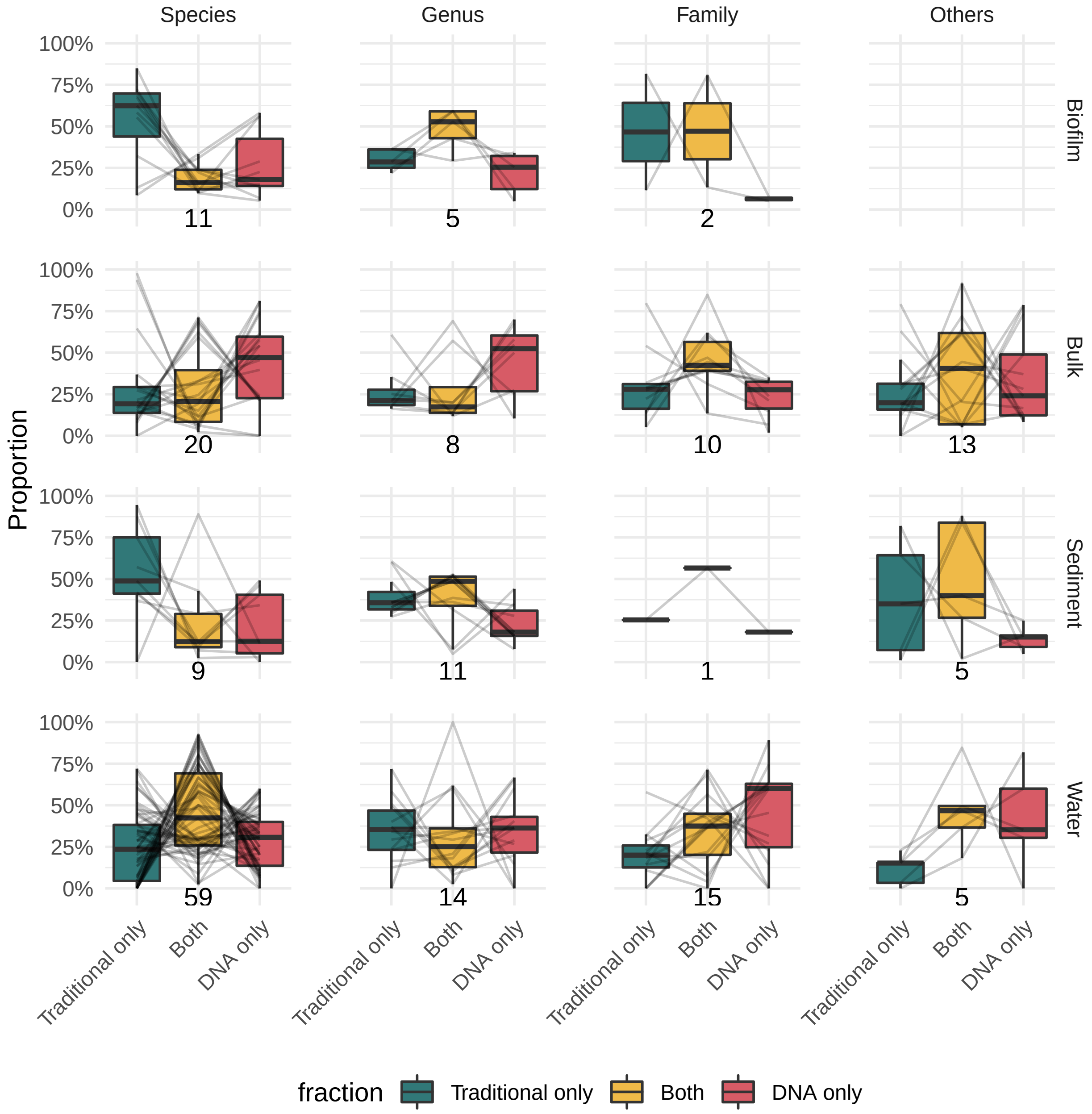

**Supplementary Figure 5.** Relative fractions of gamma diversity detected by the traditional method only, by DNA metabarcoding only and by both methods. Data are presented for different for different source of DNA and at different taxonomic levels. Boxplots show medians, first and third quartiles, and full ranges (limited to 1.5 × interquartile range). Grey lines connect values from the same comparison. Numbers below each panel indicate the number of comparisons represented.

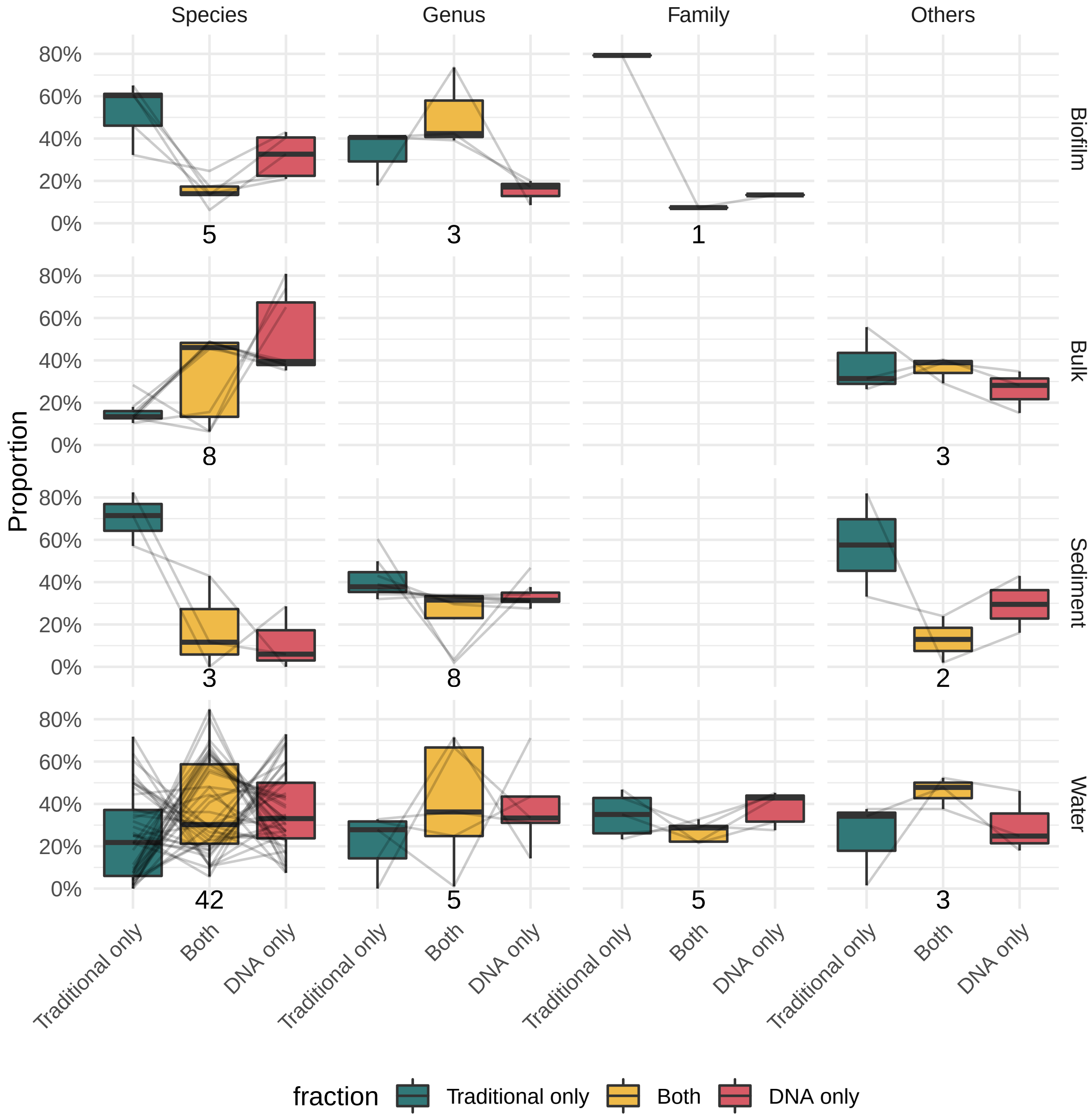

**Supplementary Figure 6.** Relative fractions of alpha diversity detected by the traditional method only, by DNA metabarcoding only and by both methods. Data are presented for different source of DNA and at different taxonomic levels. Boxplots show medians, first and third quartiles, and full ranges (limited to 1.5 × interquartile range). Grey lines connect values from the same comparison. Numbers below each panel indicate the number of comparisons represented.

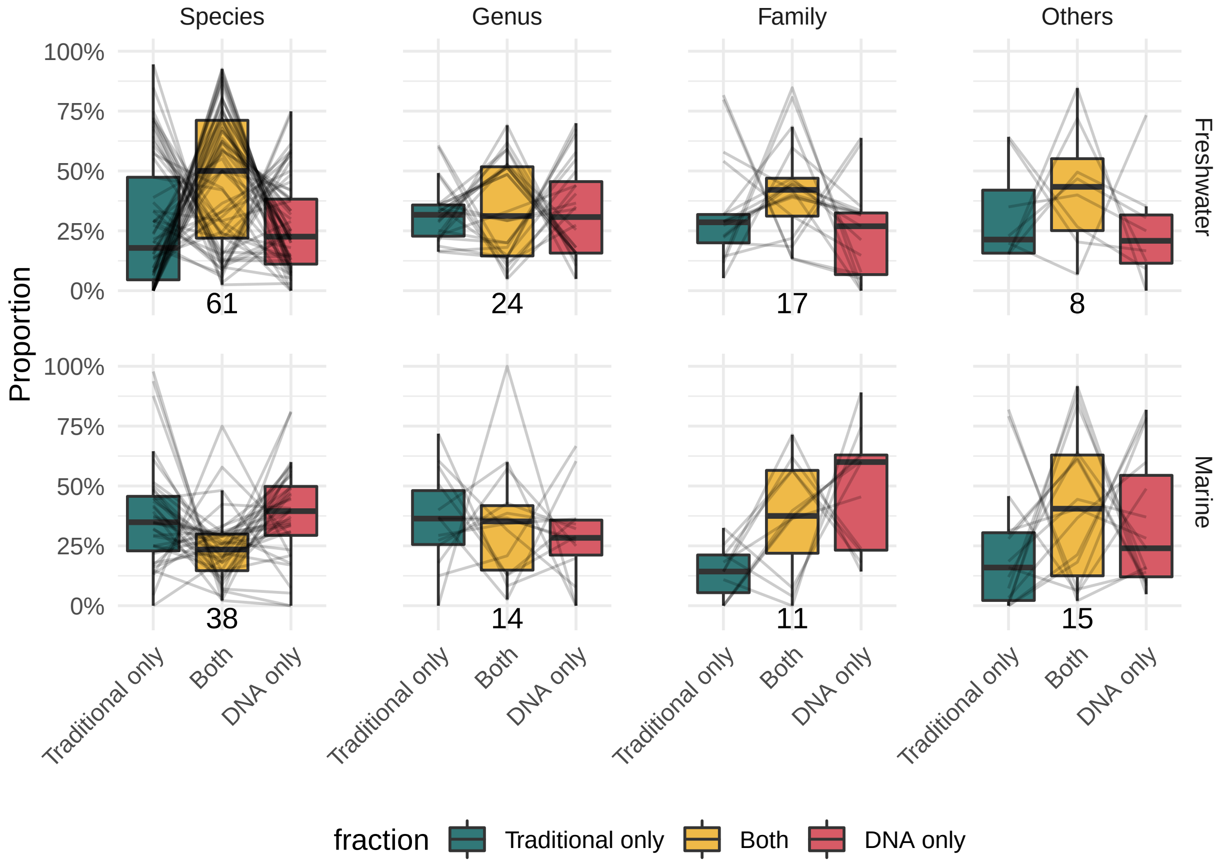

**Supplementary Figure 7.** Relative fractions of gamma diversity detected by the traditional method only, by DNA metabarcoding only and by both methods. Data are presented for different for different biomes and at different taxonomic levels. Boxplots show medians, first and third quartiles, and full ranges (limited to 1.5 × interquartile range). Grey lines connect values from the same comparison. Numbers below each panel indicate the number of comparisons represented.

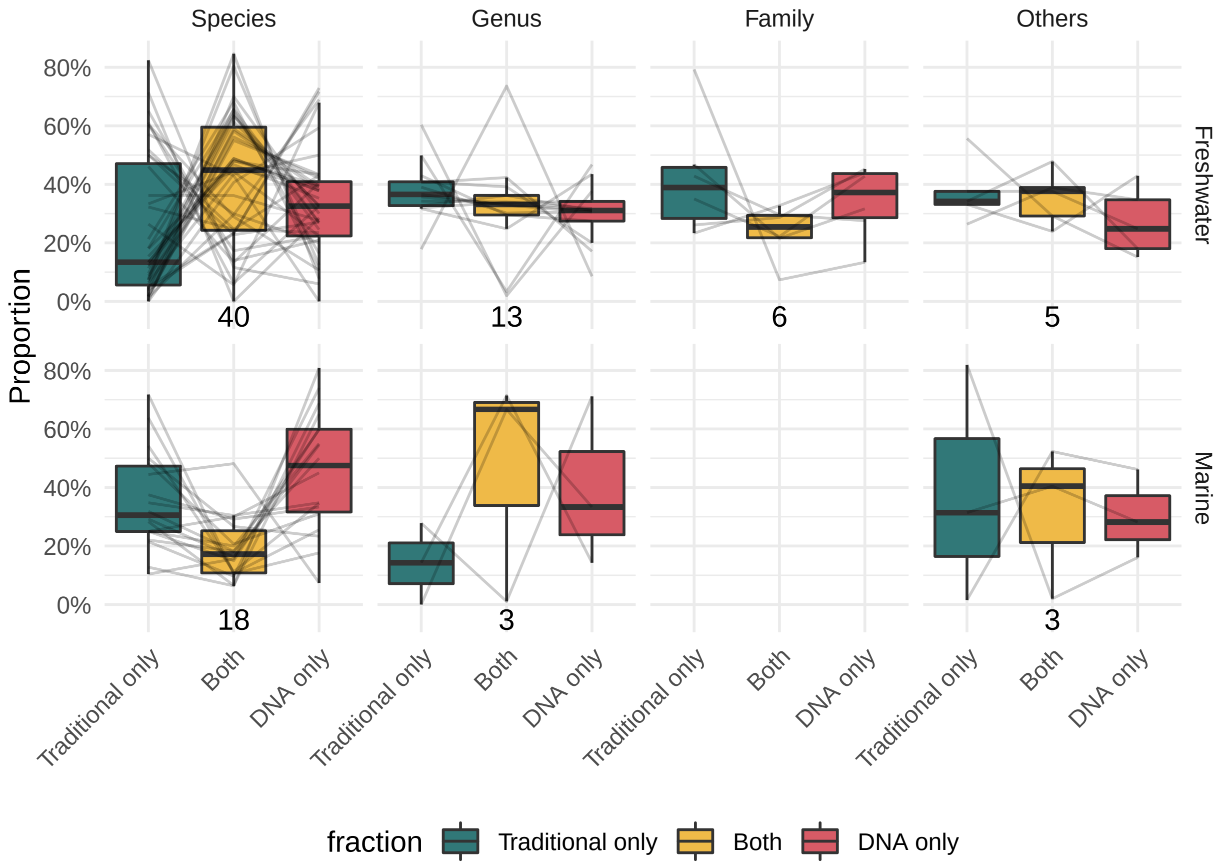

**Supplementary Figure 8.** Relative fractions of alpha diversity detected by the traditional method only, by DNA metabarcoding only and by both methods. Data are presented for different biomes and at different taxonomic levels. Boxplots show medians, first and third quartiles, and full ranges (limited to 1.5 × interquartile range). Grey lines connect values from the same comparison. Numbers below each panel indicate the number of comparisons represented.

| Variable name | Description |
|---|---|
| study_ID | ID for the study. First author name, year and first 3 letter of title. All capitals separated by underscores. |
| biome_1 | Biome level 1. |
| biome_2 | Biome level 2. |
| eDNA_origin | Origin of DNA. |
| group_target | Target of the study (PCR and traditional). |
| taxonomic_level | Taxonomic level for identification. |
| N_taxa_trad_only | Number of taxa detected only by the traditional method (in Venn diagram or in the text). |
| N_taxa_eDNA_only | Number of taxa detected only by eDNA (in Venn diagram or in the text). |
| N_taxa_both | Number of taxa detected by both methods (in Venn diagram or in the text). Intersection, not union. |
| DNA_alpha_mean | Mean richness found by DNA. |
| DNA_alpha_mean_sd | Standard deviation of the mean richness found by DNA. |
| DNA_alpha_min | Minimum richness found by DNA. |
| DNA_alpha_max | Maximum richness found by DNA. |
| trad_alpha_mean | Mean richness found by traditional method. |
| trad_alpha_mean_sd | Standard deviation of the mean richness found by traditional method. |
| trad_alpha_min | Minimum richness found by traditional method. |
| trad_alpha_max | Maximum richness found by traditional method. |
| both_alpha_mean | Mean richness found by both methods. |
| both_alpha_mean_sd | Standard deviation of the mean richness found by both methods. |
| both_alpha_min | Minimum richness found by traditional both methods. |
| both_alpha_max | Maximum richness found by traditional both methods. |
| N_sites | Number of sites represented in the comparison. |
| N_samples | Number of sample in total. |
| year_trad_sampling_start | Year traditional sampling started. |
| year_trad_sampling_end | Year traditional sampling ended. |
| year_eDNA_sampling_start | Year eDNA sampling started. |
| year_eDNA_sampling_end | Year eDNA sampling ended. |
| trad_sampling_method | Method used for traditional sampling. |
| eDNA_filtering_pore_size | Pore size for filtering eDNA (µm). |
| eDNA_filtering_volume | Volume of filtered material (ml). |
| DNA_bulk_weight | Weight of bulk material (for bulk DNA; g). |
| region_sampled | Name of the region sampled (Can be multilevel separated by column). |
| marker | Marker used for metabarcoding. |
| primer_forward | Name of the forward primer (multiprimer separated by ;). |
| primer_reverse | Name of the reverse primer (multiprimer separated by ;). |
| sequencing_technology | Name of the sequencing technology. |
| N_reads_sequencer | Total number of reads produced by the sequencer. |
| N_reads_used | Number of reads after bioinformatics and used to compute the sets. |
| N_OTU | Total number of OTUs. |
| N_OTU_target_group | Number of OTUs affiliated to the target group. |
| N_OTU_unspecific | Number of OTUS not affiliated to the target or unclassified. |
| clustering | Clustering (similarity). Use 0 for ASV/ESV. |
| refdb_coverage | Reference database used by the authors. |
| comments | Extra comments. |

**Supplementary Table 1.** List of the variables extracted from the articles.

| | Model 1 | | | Model 2 | | |
|---|---|---|---|---|---|---|
| Term | Estimate | Std. Error | P-value | Estimate | Std. Error | P-value |
| **Gamma (N = 188, Studies = 86)** | | | | | | |
| (Intercept) | -0.016 | 0.086 | 0.851 | -0.109 | 0.154 | 0.479 |
| Grp. Macroinvertebrates | | | | 0.076 | 0.214 | 0.722 |
| Grp. Fish | | | | 0.238 | 0.212 | 0.261 |
| Grp. Others | | | | -0.108 | 0.327 | 0.742 |
| **Alpha (N = 120, Studies = 61)** | | | | | | |
| (Intercept) | 0.167 | 0.085 | 0.048 | 0.180 | 0.168 | 0.286 |
| Grp. Macroinvertebrates | | | | -0.056 | 0.225 | 0.804 |
| Grp. Fish | | | | 0.042 | 0.215 | 0.845 |
| Grp. Others | | | | -0.131 | 0.309 | 0.670 |

**Supplementary Table 2.** Results of the linear mixed models explaining the log-ratio between the traditional and the metabarcoding methods for gamma and alpha diversity. Model 1 is an intercept only model. Model 2 includes the effect of the group of organisms (Grp.). Both models include the study as a random effect (intercept).

| Term | Chi-2 | Df | P-value |
|---|---|---|---|
| **Gamma** | | | |
| **(N = 291, Studies = 50, Comparisons = 97)** | | | |
| Target Group | 1.148 | 2 | 0.563 |
| Fraction | 10.732 | 2 | 0.005 |
| Target Group:Fraction | 57.360 | 4 | <0.001 |
| **Alpha** | | | |
| **(N = 168, Studies = 24, Comparisons = 56)** | | | |
| Target Group | 0.459 | 2 | 0.795 |
| Fraction | 8.234 | 2 | 0.016 |
| Target Group:Fraction | 14.069 | 4 | 0.007 |

**Supplementary Table 3.** Analysis of deviance (Type II Wald Chi-2 tests) of the generalized linear mixed models (beta regression) explaining the fractions of diversity detected by the traditional and the metabarcoding methods for gamma and alpha diversity. Models were fitted using comparisons made at species level only. Models include the effect of the type of fraction, the group of organisms (Grp.) and their interactions. Models include comparisons nested within studies as random effect (intercept).

| Paired comparisons | | | Estimate | Std. Error | P-value |
|---|---|---|---|---|---|
| **Group of organisms** | | | | | |
| Fish | Both | DNA only | 0.808 | 0.189 | <0.001 |
| | Traditional only | Both | -1.401 | 0.194 | <0.001 |
| | Traditional only | DNA only | -0.593 | 0.193 | 0.058 |
| Macroinvertebrates | Both | DNA only | -0.105 | 0.303 | >0.999 |
| | Traditional only | Both | 0.368 | 0.302 | 0.952 |
| | Traditional only | DNA only | 0.262 | 0.301 | 0.994 |
| Microorganisms | Both | DNA only | -0.537 | 0.328 | 0.783 |
| | Traditional only | Both | 1.255 | 0.327 | 0.005 |
| | Traditional only | DNA only | 0.718 | 0.322 | 0.391 |
| **Fraction** | | | | | |
| Both | Macroinvertebrates | Fish | -0.849 | 0.253 | 0.025 |
| | Microorganisms | Fish | -1.169 | 0.270 | <0.001 |
| | Microorganisms | Macroinvertebrates | -0.319 | 0.318 | 0.985 |
| DNA only | Macroinvertebrates | Fish | 0.064 | 0.252 | >0.999 |
| | Microorganisms | Fish | 0.176 | 0.265 | >0.999 |
| | Microorganisms | Macroinvertebrates | 0.112 | 0.313 | >0.999 |
| Traditional only | Macroinvertebrates | Fish | 0.920 | 0.254 | 0.010 |
| | Microorganisms | Fish | 1.487 | 0.268 | <0.001 |
| | Microorganisms | Macroinvertebrates | 0.568 | 0.310 | 0.662 |

**Supplementary Table 4.** Pairwise comparisons among factor combinations of the gamma diversity fraction model (see Supplementary Table 3) based on estimated marginal means. Only relevant comparisons (i.e. among the same group of organisms or among the same type of fraction) are displayed. P-values are adjusted using the correction of Tukey.

| Paired comparisons | | | Estimate | Std. Error | P-value |
|---|---|---|---|---|---|
| **Group of organisms** | | | | | |
| Fish | Both | DNA only | 0.120 | 0.212 | >0.999 |
| | Traditional only | Both | -0.614 | 0.215 | 0.107 |
| | Traditional only | DNA only | -0.494 | 0.215 | 0.347 |
| Macroinvertebrates | Both | DNA only | -0.664 | 0.527 | 0.941 |
| | Traditional only | Both | -0.813 | 0.534 | 0.844 |
| | Traditional only | DNA only | -1.477 | 0.537 | 0.138 |
| Microorganisms | Both | DNA only | -0.968 | 0.577 | 0.760 |
| | Traditional only | Both | 1.380 | 0.579 | 0.300 |
| | Traditional only | DNA only | 0.412 | 0.567 | 0.998 |
| **Fraction** | | | | | |
| Both | Macroinvertebrates | Fish | -0.069 | 0.401 | >0.999 |
| | Microorganisms | Fish | -0.933 | 0.442 | 0.469 |
| | Microorganisms | Macroinvertebrates | -0.865 | 0.557 | 0.829 |
| DNA only | Macroinvertebrates | Fish | 0.716 | 0.402 | 0.696 |
| | Microorganisms | Fish | 0.155 | 0.428 | >0.999 |
| | Microorganisms | Macroinvertebrates | -0.561 | 0.547 | 0.983 |
| Traditional only | Macroinvertebrates | Fish | -0.267 | 0.412 | >0.999 |
| | Microorganisms | Fish | 1.061 | 0.431 | 0.258 |
| | Microorganisms | Macroinvertebrates | 1.328 | 0.556 | 0.298 |

**Supplementary Table 5.** Pairwise comparisons among factor combinations of the alpha diversity fraction model (see Supplementary Table 3) based on estimated marginal means. Only relevant comparisons (i.e. among the same group of organisms or among the same type of fraction) are displayed. P-values are adjusted using the correction of Tukey.

**Supplementary Material 1.** Complete query used to search the Web of Science Core Collection. TS stands for topic (i.e. title, abstract and keywords), TI for title and PY for publication year. The star character (*) is a wildcard meaning zero or more character. The dollar sign ($) means zero or one character.

```
TS=(*monitor* OR *assessment* OR communit*) AND
TS=(molecular OR "environmental DNA" OR eDNA OR metabarcoding OR
    "high throughput sequencing" OR "high-throughput sequencing" OR
    HTS OR "next generation sequencing" OR "next-generation sequencing" OR NGS) AND
TS=(morpholog* OR (traditional *monitoring) OR (conventional *monitoring) OR
   (traditional *assessment*) OR (conventional *assessment*) OR "kick net" OR
   kick-net$ OR kicknet$ OR electro$fishing OR fyke$net) AND
TS=(aquatic OR freshwater$ OR marine OR stream$ OR river$ OR lake$ OR
    pond$ OR dam$ OR reservoir$ OR estuar* OR lagoon* OR sea* OR ocean$ OR coast*) AND
TS=(*invertebrate* OR Ephemeropter* OR Plecopter* OR Trichopter* OR mayfly OR
    mayflies OR caddisfl* OR stonefly OR stoneflies OR oligochaeta OR
    oligochaetes OR Diptera OR flies OR diatom* OR Bacillariophyta OR
    fish OR plant$ OR macrophyt* OR *algae) AND
PY=(2010-2025) NOT
TS=(qPCR OR ddPCR) NOT
TI=(soil OR diet OR dietary)
```

**Supplementary Material 2.** Complete list of articles included in the meta-analysis.

Abad, D., Albaina, A., Aguirre, M., Laza-Martínez, A., Uriarte, I., Iriarte, A., … Estonba, A. (2016). Is metabarcoding suitable for estuarine plankton monitoring? A comparative study with microscopy. Marine Biology, 163(7), 149. doi: 10.1007/s00227-016-2920-0

Afzali, S. F., Bourdages, H., Laporte, M., Mérot, C., Normandeau, E., Audet, C., & Bernatchez, L. (2021). Comparing environmental metabarcoding and trawling survey of demersal fish communities in the Gulf of St. Lawrence, Canada. Environmental DNA, 3(1), 22–42. doi: 10.1002/edn3.111

Aglieri, G., Baillie, C., Mariani, S., Cattano, C., Calò, A., Turco, G., … Milazzo, M. (2020). Environmental DNA effectively captures functional diversity of coastal fish communities. Molecular Ecology, mec.15661. doi: 10.1111/mec.15661

Alsos, I. G., Lammers, Y., Yoccoz, N. G., Jørgensen, T., Sjögren, P., Gielly, L., & Edwards, M. E. (2018). Plant DNA metabarcoding of lake sediments: How does it represent the contemporary vegetation. PLOS ONE, 13(4), e0195403. doi: 10.1371/journal.pone.0195403

Apothéloz-Perret-Gentil, L., Bouchez, A., Cordier, T., Cordonier, A., Guéguen, J., Rimet, F., … Pawlowski, J. (2020). Monitoring the ecological status of rivers with diatom eDNA metabarcoding: A comparison of taxonomic markers and analytical approaches for the inference of a molecular diatom index. Molecular Ecology, mec.15646. doi: 10.1111/mec.15646

Apothéloz-Perret-Gentil, L., Cordonier, A., Straub, F., Iseli, J., Esling, P., & Pawlowski, J. (2017). Taxonomy-free molecular diatom index for high-throughput eDNA biomonitoring. Molecular Ecology Resources, 17(6), 1231–1242. doi: 10.1111/1755-0998.12668

Aylagas, E., Borja, Á., Irigoien, X., & Rodríguez-Ezpeleta, N. (2016). Benchmarking DNA Metabarcoding for Biodiversity-Based Monitoring and Assessment. Frontiers in Marine Science, 3. doi: 10.3389/fmars.2016.00096

Aylagas, E., Borja, Á., Muxika, I., & Rodríguez-Ezpeleta, N. (2018). Adapting metabarcoding-based benthic biomonitoring into routine marine ecological status assessment networks. Ecological Indicators, 95, 194–202. doi: 10.1016/j.ecolind.2018.07.044

Azevedo, J., Antunes, J. T., Machado, A. M., Vasconcelos, V., Leão, P. N., & Froufe, E. (2020). Monitoring of biofouling communities in a Portuguese port using a combined morphological and metabarcoding approach. Scientific Reports, 10(1), 13461. doi: 10.1038/s41598-020-70307-4

Bachy, C., Dolan, J. R., López-García, P., Deschamps, P., & Moreira, D. (2013). Accuracy of protist diversity assessments: Morphology compared with cloning and direct pyrosequencing of 18S rRNA genes and ITS regions using the conspicuous tintinnid ciliates as a case study. The ISME Journal, 7(2), 244–255. doi: 10.1038/ismej.2012.106

Bailet, B., Apothéloz-Perret-Gentil, L., Baričević, A., Chonova, T., Franc, A., Frigerio, J.-M., … Kahlert, M. (2020). Diatom DNA metabarcoding for ecological assessment: Comparison among bioinformatics pipelines used in six European countries reveals the need for standardization. Science of The Total Environment, 745, 140948. doi: 10.1016/j.scitotenv.2020.140948

Berger, C. S., Hernandez, C., Laporte, M., Côté, G., Paradis, Y., Kameni T., D. W., … Bernatchez, L. (2020). Fine-scale environmental heterogeneity shapes fluvial fish communities as revealed by eDNA metabarcoding. Environmental DNA, 2(4), 647–666. doi: 10.1002/edn3.129

Bleijswijk, J. D. L., Engelmann, J. C., Klunder, L., Witte, H. J., Witte, J. IJ., & Veer, H. W. (2020). Analysis of a coastal North Sea fish community: Comparison of aquatic environmental DNA concentrations to fish catches. Environmental DNA, 2(4), 429–445. doi: 10.1002/edn3.67

Boivin-Delisle, D., Laporte, M., Burton, F., Dion, R., Normandeau, E., & Bernatchez, L. (2021). Using environmental DNA for biomonitoring of freshwater fish communities: Comparison with established gillnet surveys in a boreal hydroelectric impoundment. Environmental DNA, 3(1), 105–120. doi: 10.1002/edn3.135

Borrell, Y. J., Miralles, L., Do Huu, H., Mohammed-Geba, K., & Garcia-Vazquez, E. (2017). DNA in a bottle—Rapid metabarcoding survey for early alerts of invasive species in ports. PLOS ONE, 12(9), e0183347. doi: 10.1371/journal.pone.0183347

Bylemans, J., Gleeson, D. M., Lintermans, M., Hardy, C. M., Beitzel, M., Gilligan, D. M., & Furlan, E. M. (2018). Monitoring riverine fish communities through eDNA metabarcoding: Determining optimal sampling strategies along an altitudinal and biodiversity gradient. Metabarcoding and Metagenomics, 2, e30457. doi: 10.3897/mbmg.2.30457

Cahill, A. E., Pearman, J. K., Borja, A., Carugati, L., Carvalho, S., Danovaro, R., … Chenuil, A. (2018). A comparative analysis of metabarcoding and morphology-based identification of benthic communities across different regional seas. Ecology and Evolution, 8(17), 8908–8920. doi: 10.1002/ece3.4283

Cilleros, K., Valentini, A., Allard, L., Dejean, T., Etienne, R., Grenouillet, G., … Brosse, S. (2019). Unlocking biodiversity and conservation studies in high-diversity environments using environmental DNA (eDNA): A test with Guianese freshwater fishes. Molecular Ecology Resources, 19(1), 27–46. doi: 10.1111/1755-0998.12900

Clarke, L. J., Beard, J. M., Swadling, K. M., & Deagle, B. E. (2017). Effect of marker choice and thermal cycling protocol on zooplankton DNA metabarcoding studies. Ecology and Evolution, 7(3), 873–883. doi: 10.1002/ece3.2667

Closek, C. J., Santora, J. A., Starks, H. A., Schroeder, I. D., Andruszkiewicz, E. A., Sakuma, K. M., … Boehm, A. B. (2019). Marine Vertebrate Biodiversity and Distribution Within the Central California Current Using Environmental DNA (eDNA) Metabarcoding and Ecosystem Surveys. Frontiers in Marine Science, 6, 732. doi: 10.3389/fmars.2019.00732

Collins, R. A., Bakker, J., Wangensteen, O. S., Soto, A. Z., Corrigan, L., Sims, D. W., … Mariani, S. (2019). Non-specific amplification compromises environmental DNA metabarcoding with COI. Methods in Ecology and Evolution, 10(11), 1985–2001. doi: 10.1111/2041-210X.13276

Cowart, D. A., Pinheiro, M., Mouchel, O., Maguer, M., Grall, J., Miné, J., & Arnaud-Haond, S. (2015). Metabarcoding Is Powerful yet Still Blind: A Comparative Analysis of Morphological and Molecular Surveys of Seagrass Communities. PLOS ONE, 10(2), e0117562. doi: 10.1371/journal.pone.0117562

Deagle, B. E., Clarke, L. J., Kitchener, J. A., Polanowski, A. M., & Davidson, A. T. (2018). Genetic monitoring of open ocean biodiversity: An evaluation of DNA metabarcoding for processing continuous plankton recorder samples. Molecular Ecology Resources, 18(3), 391–406. doi: 10.1111/1755-0998.12740

Djurhuus, A., Pitz, K., Sawaya, N. A., Rojas-Márquez, J., Michaud, B., Montes, E., … Breitbart, M. (2018). Evaluation of marine zooplankton community structure through environmental DNA metabarcoding: Metabarcoding zooplankton from eDNA. Limnology and Oceanography: Methods, 16(4), 209–221. doi: 10.1002/lom3.10237

Doble, C. J., Hipperson, H., Salzburger, W., Horsburgh, G. J., Mwita, C., Murrell, D. J., & Day, J. J. (2020). Testing the performance of environmental DNA metabarcoding for surveying highly diverse tropical fish communities: A case study from Lake Tanganyika. Environmental DNA, 2(1), 24–41. doi: 10.1002/edn3.43

Dzhembekova, N., Rubino, F., Nagai, S., Zlateva, I., Slabakova, N., Ivanova, P., … Moncheva, S. (2020). Comparative analysis of morphological and molecular approaches integrated into the study of the dinoflagellate biodiversity within the recently deposited Black Sea sediments – benefits and drawbacks. Biodiversity Data Journal, 8, e55172. doi: 10.3897/BDJ.8.e55172

Eiler, A., Drakare, S., Bertilsson, S., Pernthaler, J., Peura, S., Rofner, C., … Lindström, E. S. (2013). Unveiling Distribution Patterns of Freshwater Phytoplankton by a Next Generation Sequencing Based Approach. PLoS ONE, 8(1), e53516. doi: 10.1371/journal.pone.0053516

Elbrecht, V., Vamos, E. E., Meissner, K., Aroviita, J., & Leese, F. (2017). Assessing strengths and weaknesses of DNA metabarcoding-based macroinvertebrate identification for routine stream monitoring. Methods in

Ecology and Evolution, 8(10), 1265–1275. doi: 10.1111/2041-210X.12789

Emilson, C. E., Thompson, D. G., Venier, L. A., Porter, T. M., Swystun, T., Chartrand, D., … Hajibabaei, M. (2017). DNA metabarcoding and morphological macroinvertebrate metrics reveal the same changes in boreal watersheds across an environmental gradient. Scientific Reports, 7(1), 12777. doi: 10.1038/s41598-017-13157-x

Erdozain, M., Thompson, D. G., Porter, T. M., Kidd, K. A., Kreutzweiser, D. P., Sibley, P. K., … Hajibabaei, M. (2019). Metabarcoding of storage ethanol vs. Conventional morphometric identification in relation to the use of stream macroinvertebrates as ecological indicators in forest management. Ecological Indicators, 101, 173–184. doi: 10.1016/j.ecolind.2019.01.014

Evans, N. T., Li, Y., Renshaw, M. A., Olds, B. P., Deiner, K., Turner, C. R., … Pfrender, M. E. (2017). Fish community assessment with eDNA metabarcoding: Effects of sampling design and bioinformatic filtering. Canadian Journal of Fisheries and Aquatic Sciences, 74(9), 1362–1374. doi: 10.1139/cjfas-2016-0306

Fernández, S., Rodríguez, S., Martínez, J. L., Borrell, Y. J., Ardura, A., & García-Vázquez, E. (2018). Evaluating freshwater macroinvertebrates from eDNA metabarcoding: A river Nalón case study. PLOS ONE, 13(8), e0201741. doi: 10.1371/journal.pone.0201741

Fernández, S., Rodríguez-Martínez, S., Martínez, J. L., Garcia-Vazquez, E., & Ardura, A. (2019). How can eDNA contribute in riverine macroinvertebrate assessment? A metabarcoding approach in the Nalón River (Asturias, Northern Spain). Environmental DNA, 1(4), 385–401. doi: 10.1002/edn3.40

Fujii, K., Doi, H., Matsuoka, S., Nagano, M., Sato, H., & Yamanaka, H. (2019). Environmental DNA metabarcoding for fish community analysis in backwater lakes: A comparison of capture methods. PLOS ONE, 14(1), e0210357. doi: 10.1371/journal.pone.0210357

Goutte, A., Molbert, N., Guérin, S., Richoux, R., & Rocher, V. (2020). Monitoring freshwater fish communities in large rivers using environmental DNA metabarcoding and a long-term electrofishing survey. Journal of Fish Biology, 97(2), 444–452. doi: 10.1111/jfb.14383

Gran-Stadniczeñko, S., Šupraha, L., Egge, E. D., & Edvardsen, B. (2017). Haptophyte Diversity and Vertical Distribution Explored by 18S and 28S Ribosomal RNA Gene Metabarcoding and Scanning Electron Microscopy. Journal of Eukaryotic Microbiology, 64(4), 514–532. doi: 10.1111/jeu.12388

Haenel, Q., Holovachov, O., Jondelius, U., Sundberg, P., & Bourlat, S. (2017). NGS-based biodiversity and community structure analysis of meiofaunal eukaryotes in shell sand from Hållö island, Smögen, and soft mud from Gullmarn Fjord, Sweden. Biodiversity Data Journal, 5, e12731. doi: 10.3897/BDJ.5.e12731

Hänfling, B., Lawson Handley, L., Read, D. S., Hahn, C., Li, J., Nichols, P., … Winfield, I. J. (2016). Environmental DNA metabarcoding of lake fish communities reflects long-term data from established survey methods. Molecular Ecology, 25(13), 3101–3119. doi: 10.1111/mec.13660

Harper, L. R., Lawson Handley, L., Sayer, C. D., Read, D. S., Benucci, M., Blackman, R. C., … Hänfling, B. (2020). Assessing the impact of the threatened crucian carp ( Carassius carassius ) on pond invertebrate diversity: A comparison of conventional and molecular tools. Molecular Ecology, mec.15670. doi: 10.1111/mec.15670

Harvey, J. B. J., Johnson, S. B., Fisher, J. L., Peterson, W. T., & Vrijenhoek, R. C. (2017). Comparison of morphological and next generation DNA sequencing methods for assessing zooplankton assemblages. Journal of Experimental Marine Biology and Ecology, 487, 113–126. doi: 10.1016/j.jembe.2016.12.002

Hayami, K., Sakata, M. K., Inagawa, T., Okitsu, J., Katano, I., Doi, H., … Minamoto, T. (2020). Effects of sampling seasons and locations on fish environmental DNA metabarcoding in dam reservoirs. Ecology and Evolution, 10(12), 5354–5367. doi: 10.1002/ece3.6279

Hirai, J., Yasuike, M., Fujiwara, A., Nakamura, Y., Hamaoka, S., Katakura, S., … Nagai, S. (2015). Effects of plankton net characteristics on metagenetic community analysis of metazoan zooplankton in a coastal marine ecosystem. Journal of Experimental Marine Biology and Ecology, 469, 36–43. doi: 10.1016/j.jembe.2015.04.011

Huang, S., Herzschuh, U., Pestryakova, L. A., Zimmermann, H. H., Davydova, P., Biskaborn, B. K., … Stoof-Leichsenring, K. R. (2020). Genetic and morphologic determination of diatom community composition in surface sediments from glacial and thermokarst lakes in the Siberian Arctic. Journal of Paleolimnology, 64(3), 225–242. doi: 10.1007/s10933-020-00133-1

Huo, S., Li, X., Xi, B., Zhang, H., Ma, C., & He, Z. (2020). Combining morphological and metabarcoding approaches reveals the freshwater eukaryotic phytoplankton community. Environmental Sciences Europe, 32(1), 37. doi: 10.1186/s12302-020-00321-w

Kang, W., Anslan, S., Börner, N., Schwarz, A., Schmidt, R., Künzel, S., … Schwalb, A. (2021). Diatom metabarcoding and microscopic analyses from sediment samples at Lake Nam Co, Tibet: The effect of sample-size and bioinformatics on the identified communities. Ecological Indicators, 121, 107070. doi: 10.1016/j.ecolind.2020.107070

Kelly, R. P., Closek, C. J., O'Donnell, J. L., Kralj, J. E., Shelton, A. O., & Samhouri, J. F. (2017). Genetic and Manual Survey Methods Yield Different and Complementary Views of an Ecosystem. Frontiers in Marine Science, 3. doi: 10.3389/fmars.2016.00283

Kermarrec, L., Franc, A., Rimet, F., Chaumeil, P., Frigerio, J.-M., Humbert, J.-F., & Bouchez, A. (2014). A next-generation sequencing approach to river biomonitoring using benthic diatoms. Freshwater Science, 33(1), 349–363. doi: 10.1086/675079

Kim, H., Lee, C.-R., Lee, S., Oh, S.-Y., & Kim, W. (2020). Biodiversity and Community Structure of Mesozooplankton in the Marine and Coastal National Park Areas of Korea. Diversity, 12(6), 233. doi: 10.3390/d12060233

Kim, Park, Jo, & Kwak. (2019). Comparison of Water Sampling between Environmental DNA Metabarcoding and Conventional Microscopic Identification: A Case Study in Gwangyang Bay, South Korea. Applied Sciences, 9(16), 3272. doi: 10.3390/app9163272

Krol, L., Van der Hoorn, B., Gorsich, E. E., Trimbos, K., Bodegom, P. M. van, & Schrama, M. (2019). How Does eDNA Compare to Traditional Trapping? Detecting Mosquito Communities in South-African Freshwater Ponds. Frontiers in Ecology and Evolution, 7, 260. doi: 10.3389/fevo.2019.00260

Kuntke, F., de Jonge, N., Hesselsøe, M., & Lund Nielsen, J. (2020). Stream water quality assessment by metabarcoding of invertebrates. Ecological Indicators, 111, 105982. doi: 10.1016/j.ecolind.2019.105982

Laini, A., Beermann, A. J., Bolpagni, R., Burgazzi, G., Elbrecht, V., Zizka, V. M. A., … Viaroli, P. (2020). Exploring the potential of metabarcoding to disentangle macroinvertebrate community dynamics in intermittent streams. Metabarcoding and Metagenomics, 4, e51433. doi: 10.3897/mbmg.4.51433

Leduc, N., Lacoursière-Roussel, A., Howland, K. L., Archambault, P., Sevellec, M., Normandeau, E., … Bernatchez, L. (2019). Comparing eDNA metabarcoding and species collection for documenting Arctic metazoan biodiversity. Environmental DNA, 1(4), 342–358. doi: 10.1002/edn3.35

Leese, F., Sander, M., Buchner, D., Elbrecht, V., Haase, P., & Zizka, V. M. A. (2021). Improved freshwater macroinvertebrate detection from environmental DNA through minimized nontarget amplification. Environmental DNA, 3(1), 261–276. doi: 10.1002/edn3.177

Lejzerowicz, F., Esling, P., Pillet, L., Wilding, T. A., Black, K. D., & Pawlowski, J. (2015). High-throughput sequencing and morphology perform equally well for benthic monitoring of marine ecosystems. Scientific Reports, 5(1), 13932. doi: 10.1038/srep13932

Li, X., Huo, S., Zhang, J., Ma, C., Xiao, Z., Zhang, H., … Xia, X. (2019). Metabarcoding reveals a more complex cyanobacterial community than morphological identification. Ecological Indicators, 107, 105653. doi: 10.1016/j.ecolind.2019.105653

Liu, Y., Song, S., Chen, T., & Li, C. (2017). The diversity and structure of marine protists in the coastal waters of China revealed by morphological observation and 454 pyrosequencing. Estuarine, Coastal and Shelf Science, 189, 143–155. doi: 10.1016/j.ecss.2017.03.019

Lobo, J., Shokralla, S., Costa, M. H., Hajibabaei, M., & Costa, F. O. (2017). DNA metabarcoding for high-throughput monitoring of estuarine macrobenthic communities. Scientific Reports, 7(1), 15618. doi: 10.1038/s41598-017-15823-6

Mächler, E., Little, C. J., Wüthrich, R., Alther, R., Fronhofer, E. A., Gounand, I., … Altermatt, F. (2019). Assessing different components of diversity across a river network using eDNA. Environmental DNA, 1(3), 290–301. doi: 10.1002/edn3.33

Marshall, N. T., & Stepien, C. A. (2020). Macroinvertebrate community diversity and habitat quality relationships along a large river from targeted eDNA metabarcode assays. Environmental DNA, 2(4), 572–586. doi: 10.1002/edn3.90

Martins, F. M. S., Galhardo, M., Filipe, A. F., Teixeira, A., Pinheiro, P., Paupério, J., … Beja, P. (2019). Have the cake and eat it: Optimizing nondestructive DNA metabarcoding of macroinvertebrate samples for freshwater biomonitoring. Molecular Ecology Resources, 19(4), 863–876. doi: 10.1111/1755-0998.13012

McClenaghan, B., Fahner, N., Cote, D., Chawarski, J., McCarthy, A., Rajabi, H., … Hajibabaei, M. (2020). Harnessing the power of eDNA metabarcoding for the detection of deep-sea fishes. PLOS ONE, 15(11), e0236540. doi: 10.1371/journal.pone.0236540

McDevitt, A. D., Sales, N. G., Browett, S. S., Sparnenn, A. O., Mariani, S., Wangensteen, O. S., … Benvenuto, C. (2019). Environmental DNA metabarcoding as an effective and rapid tool for fish monitoring in canals. Journal of Fish Biology, 95(2), 679–682. doi: 10.1111/jfb.14053

Minerovic, A. D., Potapova, M. G., Sales, C. M., Price, J. R., & Enache, M. D. (2020). 18S-V9 DNA metabarcoding detects the effect of water-quality impairment on stream biofilm eukaryotic assemblages. Ecological Indicators, 113, 106225. doi: 10.1016/j.ecolind.2020.106225

Mora, D., Abarca, N., Proft, S., Grau, J. H., Enke, N., Carmona, J., … Zimmermann, J. (2019). Morphology and metabarcoding: A test with stream diatoms from Mexico highlights the complementarity of identification methods. Freshwater Science, 38(3), 448–464. doi: 10.1086/704827

Nguyen, B. N., Shen, E. W., Seemann, J., Correa, A. M. S., O'Donnell, J. L., Altieri, A. H., … Leray, M. (2020). Environmental DNA survey captures patterns of fish and invertebrate diversity across a tropical seascape. Scientific Reports, 10(1), 6729. doi: 10.1038/s41598-020-63565-9

Nichols, P. K., & Marko, P. B. (2019). Rapid assessment of coral cover from environmental DNA in Hawai'i. Environmental DNA, 1(1), 40–53. doi: 10.1002/edn3.8

Nunes, M., Adams, J., Van Aswegen, S., & Matcher, G. (2019). A comparison between the morphological and molecular approach to identify the benthic diatom community in the St Lucia Estuary, South Africa. African Journal of Marine Science, 41(4), 429–442. doi: 10.2989/1814232X.2019.1689169

Obst, M., Exter, K., Allcock, A. L., Arvanitidis, C., Axberg, A., Bustamante, M., … Pavloudi, C. (2020). A Marine Biodiversity Observation Network for Genetic Monitoring of Hard-Bottom Communities (ARMS-MBON). Frontiers in Marine Science, 7, 572680. doi: 10.3389/fmars.2020.572680

Oka, S., Doi, H., Miyamoto, K., Hanahara, N., Sado, T., & Miya, M. (2021). Environmental DNA metabarcoding for biodiversity monitoring of a highly diverse tropical fish community in a coral reef lagoon: Estimation of species richness and detection of habitat segregation. Environmental DNA, 3(1), 55–69. doi: 10.1002/edn3.132

Olds, B. P., Jerde, C. L., Renshaw, M. A., Li, Y., Evans, N. T., Turner, C. R., … Lamberti, G. A. (2016). Estimating species richness using environmental DNA. Ecology and Evolution, 6(12), 4214–4226. doi: 10.1002/ece3.2186

Pérez-Burillo, J., Trobajo, R., Vasselon, V., Rimet, F., Bouchez, A., & Mann, D. G. (2020). Evaluation and sensitivity analysis of diatom DNA metabarcoding for WFD bioassessment of Mediterranean rivers. Science of The Total Environment, 727, 138445. doi: 10.1016/j.scitotenv.2020.138445

Polanco Fernández, A., Marques, V., Fopp, F., Juhel, J., Borrero-Pérez, G. H., Cheutin, M., … Pellissier, L. (2021). Comparing environmental DNA metabarcoding and underwater visual census to monitor tropical reef fishes. Environmental DNA, 3(1), 142–156. doi: 10.1002/edn3.140

Pont, D., Rocle, M., Valentini, A., Civade, R., Jean, P., Maire, A., … Dejean, T. (2018). Environmental DNA reveals quantitative patterns of fish biodiversity in large rivers despite its downstream transportation. Scientific Reports, 8(1), 10361. doi: 10.1038/s41598-018-28424-8

Port, J. A., O'Donnell, J. L., Romero-Maraccini, O. C., Leary, P. R., Litvin, S. Y., Nickols, K. J., … Kelly, R. P. (2016). Assessing vertebrate biodiversity in a kelp forest ecosystem using environmental DNA. Molecular Ecology, 25(2), 527–541. doi: 10.1111/mec.13481

Pujari, L., Wu, C., Kan, J., Li, N., Wang, X., Zhang, G., … Sun, J. (2019). Diversity and Spatial Distribution of Chromophytic Phytoplankton in the Bay of Bengal Revealed by RuBisCO Genes (rbcL). Frontiers in Microbiology, 10, 1501. doi: 10.3389/fmicb.2019.01501

Rivera, S. F., Vasselon, V., Jacquet, S., Bouchez, A., Ariztegui, D., & Rimet, F. (2018). Metabarcoding of lake benthic diatoms: From structure assemblages to ecological assessment. Hydrobiologia, 807(1), 37–51. doi: 10.1007/s10750-017-3381-2

Rivera, Sinziana F., Vasselon, V., Ballorain, K., Carpentier, A., Wetzel, C. E., Ector, L., … Rimet, F. (2018). DNA metabarcoding and microscopic analyses of sea turtles biofilms: Complementary to understand turtle behavior. PLOS ONE, 13(4), e0195770. doi: 10.1371/journal.pone.0195770

Rivera, Sinziana F., Vasselon, V., Mary, N., Monnier, O., Rimet, F., & Bouchez, A. (2021). Exploring the capacity of aquatic biofilms to act as environmental DNA samplers: Test on macroinvertebrate communities in rivers. Science of The Total Environment, 763, 144208. doi: 10.1016/j.scitotenv.2020.144208

Sakata, M. K., Watanabe, T., Maki, N., Ikeda, K., Kosuge, T., Okada, H., … Minamoto, T. (2021). Determining an effective sampling method for eDNA metabarcoding: A case study for fish biodiversity monitoring in a small, natural river. Limnology, 22(2), 221–235. doi: 10.1007/s10201-020-00645-9

Sard, N. M., Herbst, S. J., Nathan, L., Uhrig, G., Kanefsky, J., Robinson, J. D., & Scribner, K. T. (2019). Comparison of fish detections, community diversity, and relative abundance using environmental DNA metabarcoding and traditional gears. Environmental DNA, 1(4), 368–384. doi: 10.1002/edn3.38

Schroeder, A., Stanković, D., Pallavicini, A., Gionechetti, F., Pansera, M., & Camatti, E. (2020). DNA metabarcoding and morphological analysis—Assessment of zooplankton biodiversity in transitional waters. Marine Environmental Research, 160, 104946. doi: 10.1016/j.marenvres.2020.104946

Semmouri, I., De Schamphelaere, K. A. C., Willemse, S., Vandegehuchte, M. B., Janssen, C. R., & Asselman, J. (2021). Metabarcoding reveals hidden species and improves identification of marine zooplankton communities in the North Sea. ICES Journal of Marine Science, fsaa256. doi: 10.1093/icesjms/fsaa256

Serrana, J. M., Miyake, Y., Gamboa, M., & Watanabe, K. (2019). Comparison of DNA metabarcoding and morphological identification for stream macroinvertebrate biodiversity assessment and monitoring. Ecological Indicators, 101, 963–972. doi: 10.1016/j.ecolind.2019.02.008

Serrana, J. M., Yaegashi, S., Kondoh, S., Li, B., Robinson, C. T., & Watanabe, K. (2018). Ecological influence of sediment bypass tunnels on macroinvertebrates in dam-fragmented rivers by DNA metabarcoding. Scientific Reports, 8(1), 10185. doi: 10.1038/s41598-018-28624-2

Shackleton, M. E., Rees, G. N., Watson, G., Campbell, C., & Nielsen, D. (2019). Environmental DNA reveals landscape mosaic of wetland plant communities. Global Ecology and Conservation, 19, e00689. doi: 10.1016/j.gecco.2019.e00689

Shaw, J. L. A., Clarke, L. J., Wedderburn, S. D., Barnes, T. C., Weyrich, L. S., & Cooper, A. (2016). Comparison of environmental DNA metabarcoding and conventional fish survey methods in a river system. Biological Conservation, 197, 131–138. doi: 10.1016/j.biocon.2016.03.010

Snyder, M. R., & Stepien, C. A. (2020). Increasing confidence for discerning species and population compositions from metabarcoding assays of environmental samples: Case studies of fishes in the Laurentian Great Lakes and Wabash River. Metabarcoding and Metagenomics, 4, e53455. doi: 10.3897/mbmg.4.53455

Steyaert, M., Priestley, V., Osborne, O., Herraiz, A., Arnold, R., & Savolainen, V. (2020). Advances in metabarcoding techniques bring us closer to reliable monitoring of the marine benthos. Journal of Applied Ecology, 57(11), 2234–2245. doi: 10.1111/1365-2664.13729

Sun, Z., Majaneva, M., Sokolova, E., Rauch, S., Meland, S., & Ekrem, T. (2019). DNA metabarcoding adds valuable information for management of biodiversity in roadside stormwater ponds. Ecology and Evolution, 9(17), 9712–9722. doi: 10.1002/ece3.5503

Thomsen, P. F., Kielgast, J., Iversen, L. L., Møller, P. R., Rasmussen, M., & Willerslev, E. (2012). Detection of a Diverse Marine Fish Fauna Using Environmental DNA from Seawater Samples. PLoS ONE, 7(8), e41732. doi: 10.1371/journal.pone.0041732

Uchida, N., Kubota, K., Aita, S., & Kazama, S. (2020). Aquatic insect community structure revealed by eDNA metabarcoding derives indices for environmental assessment. PeerJ, 8, e9176. doi: 10.7717/peerj.9176

Valentini, A., Taberlet, P., Miaud, C., Civade, R., Herder, J., Thomsen, P. F., … Dejean, T. (2016). Next-generation monitoring of aquatic biodiversity using environmental DNA metabarcoding. Molecular Ecology, 25(4), 929–942. doi: 10.1111/mec.13428

Vasselon, V., Rimet, F., Tapolczai, K., & Bouchez, A. (2017). Assessing ecological status with diatoms DNA metabarcoding: Scaling-up on a WFD monitoring network (Mayotte island, France). Ecological Indicators, 82, 1–12. doi: 10.1016/j.ecolind.2017.06.024

Visco, J. A., Apothéloz-Perret-Gentil, L., Cordonier, A., Esling, P., Pillet, L., & Pawlowski, J. (2015). Environmental Monitoring: Inferring the Diatom Index from Next-Generation Sequencing Data. Environmental Science & Technology, 49(13), 7597–7605. doi: 10.1021/es506158m

Vivien, R., Apothéloz-Perret-Gentil, L., Pawlowski, J., Werner, I., & Ferrari, B. J. D. (2019). Testing different (e)DNA metabarcoding approaches to assess aquatic oligochaete diversity and the biological quality of sediments. Ecological Indicators, 106, 105453. doi: 10.1016/j.ecolind.2019.105453

Xiao, X., Sogge, H., Lagesen, K., Tooming-Klunderud, A., Jakobsen, K. S., & Rohrlack, T. (2014). Use of High Throughput Sequencing and Light Microscopy Show Contrasting Results in a Study of Phytoplankton Occurrence in a Freshwater Environment. PLoS ONE, 9(8), e106510. doi: 10.1371/journal.pone.0106510

Yang, J., Zhang, X., Xie, Y., Song, C., Zhang, Y., Yu, H., & Burton, G. A. (2017). Zooplankton Community Profiling in a Eutrophic Freshwater Ecosystem-Lake Tai Basin by DNA Metabarcoding. Scientific Reports, 7(1), 1773. doi: 10.1038/s41598-017-01808-y

Zimmermann, J., Glöckner, G., Jahn, R., Enke, N., & Gemeinholzer, B. (2015). Metabarcoding vs. Morphological identification to assess diatom diversity in environmental studies. Molecular Ecology Resources, 15(3), 526–542. doi: 10.1111/1755-0998.12336
